# Primary auditory cortex represents the location of sound sources in a cue- invariant manner

**DOI:** 10.1101/348672

**Authors:** Katherine C Wood, Stephen M Town, Jennifer K Bizley

## Abstract

Auditory cortex is required for sound localisation, but how neural firing in auditory cortex underlies our perception of sources in space remains unknown. We measured spatial receptive fields in animals actively attending to spatial location while they performed a relative localisation task using stimuli that varied in the spatial cues that they provided. Manipulating the availability of binaural and spectral localisation cues had mild effects on the ferret’s performance and little impact on the spatial tuning of neurons in primary auditory cortex (A1). Consistent with a representation of space, a subpopulation of neurons encoded spatial position across localisation cue types. Spatial receptive fields measured in the presence of a competing sound source were sharper than those measured in a single-source configuration. Together these observations suggest that A1 encodes the location of auditory objects as opposed to spatial cue values. We compared our data to predictions generated from two theories about how space is represented in auditory cortex: The two-channel model, where location is encoded by the relative activity in each hemisphere, and the labelled-line model where location is represented by the activity pattern of individual cells. The representation of sound location in A1 was mainly contralateral but peak firing rates were distributed across the hemifield consistent with a labelled line model in each hemisphere representing contralateral space. Comparing reconstructions of sound location from neural activity, we found that a labelled line architecture far outperformed two channel systems. Reconstruction ability increased with increasing channel number, saturating at around 20 channels.

**Significance statement:** Our perception of a sound scene is one of distinct sound sources each of which can be localised, yet auditory space must be computed from sound location cues that arise principally by comparing the sound at the two ears. Here we ask: (1) do individual neurons in auditory cortex represent space, or sound localisation cues? (2) How is neural activity ‘read out’ for spatial perception? We recorded from auditory cortex in ferrets performing a localisation task and describe a subpopulation of neurons that represent space across localisation cues. Our data are consistent with auditory space being read out using the pattern of activity across neurons (a labelled line) rather than by averaging activity within each hemisphere (a two-channel model).

## Introduction

Our ability to localise sounds is important for both survival and communication. Auditory cortex is required for sound localisation in many mammals, including primates, cats and ferrets (1–5), and neurons in auditory cortex are tuned to the spatial location of sounds (e.g. Middlebrooks and Pettigrew, 1981; Recanzone et al., 2000; Town et al., 2017). However, perceptual thresholds for localisation are typically far narrower than tuning of auditory cortical neurons (6, 13–15), posing the question of how such cortical activity is related to spatial perception.

Studies of spatial representations in auditory cortex have often focussed on the encoding of the acoustic cues that support sound localisation. These include binaural cues such as inter-aural timing and level differences (ITDs and ILDs respectively) that govern estimations of azimuthal sound location (Middlebrooks and Green, 1991; Keating et al., 2014), as well as spectral cues arising from pinna shape (8, 19–21). While spatial cues can provide redundant information, accurate sound localisation in multisource or reverberant environments frequently requires that we integrate information across more than one cue type (22, 23). Indeed binaural and spectral cues are combined within the inferior colliculus (24, 25), potentially enabling cortical representations that are better characterized by sound location than particular acoustic features (26, 27). While it is clear that most (but not all) neurons in A1 represent the location of a sound source relative to the head (i.e. an egocentric representation, potentially indicative of a cue-based representation (11)), it remains unclear whether neurons in auditory cortex represent localisation cues or a more integrated representation of ‘space’ since most studies do not consider the effects of systematically manipulating the available localisation cues on coding of spatial location.

Sound location plays a critical role in the analysis of auditory scenes and the formation of auditory objects: perceptual representations of a sound source in the world which are both identifiable and perceived as originating from a distinct location (28, 29). Auditory cortex has been proposed to have a key role in the formation of auditory objects (30). Object based representations are often elucidated by using stimulus competition: For example, two sounds that are presented from different locations can be fused together as a single perceptual source if other features of the sound imply that the sounds are from the same sound source (31). In contrast, when two competing sources are experienced as distinct objects, they can ‘repel’, causing human listeners to perceive them as further apart than they are in reality (32). Consistent with an object based representation, in auditory cortex, the presence of a competing sound source causes dramatic enhancement of spatial tuning properties in auditory cortical neurons (33, 34) and when two sources are perceived as perceptually fused, auditory cortical activity is consistent with the location of the fused percept, i.e. between the two physical sound sources (35).

Although it is unclear *what* auditory cortical neurons are representing with regard to sound location, several models exist of *how* neurons and neural populations represent sound location: The labelled line model (17, 36) posits that cells have heterogeneous spatial tuning to many sound locations (or their underlying acoustic cues) whereas the two-channel model (37, 38) suggests that tuning is broad and conserved across cells within a hemisphere with space represented by the relative summed activity of each hemisphere.

There is both supporting and conflicting evidence for each potential mechanism of spatial representation. In agreement with the two channel model, spatial tuning curves of neurons in auditory cortex are generally broadly tuned with peaks in contralateral space (10, 11, 37) and functional imaging data is consistent with a two-channel representation (26, 38). In contrast, and consistent with the labelled-line hypothesis, neurons in a single hemisphere of gerbil auditory cortex were found to represent ITDs across all sound locations rather than just a single side of space (39). However both a two-channel representation, and a labelled line model in which each hemisphere contains a panoramic representation of the azimuthal plane cannot account for the contralateral localization deficit observed with unilateral inactivation of auditory cortex (13, 14, 40). To resolve the neural code for sound space, it is necessary to record neural activity with high spatial and temporal precision to avoid averaging signals over large populations of neurons (which may artificially favour the two channel model). Here we achieved this using extracellular electrophysiology in animals discriminating sound location.

The goal of this study was to address the question of whether neurons in auditory cortex represent spatial cue values or sound source location, and to determine how population activity represents sound source location. Since behavioural state can alter spatial tuning (12) recordings were made while animals were discriminating the azimuthal location of sounds which varied in the availability of localisation cues. The resulting spatial receptive fields of neurons were used to assess whether AC encodes ‘space’ or auditory cue values, and whether a labelled line or two-channel model provides the best description of the observed data. We hypothesised that if auditory cortex encoded space, rather than spatial cues, then spatial receptive fields should be invariant to available localisation cues. We found that spatial receptive fields were largely robust across variation in cue availability and type, consistent with the representation of spatial location. Consistent with a neural representation of the location of a source, a competing stimulus sharpened spatial receptive fields. We also observed that the best azimuths of neurons in A1 were distributed across the contralateral hemifield tiling space in a labelled-line manner and that a labelled-line population decoder outperformed a two-channel decoder.

## Results

### Relative localization judgments using complete and cue restricted stimuli

To understand the representation of space in auditory cortex, we engaged animals in a task that required attention to the azimuthal location of a sound source. Ferrets performed a two-interval forced choice task in which they reported whether a target sound was presented to the left or right of a preceding reference (Fig. 1). Reference stimuli were presented from −75° to +75° in azimuth in 30° steps at 0° elevation, with subsequent target sounds occurring 30° to the left or right of the reference location (at ± 75°, targets always moved towards the midline). Both reference and target sounds were 150 ms duration, separated by 20 ms of silence. Acoustic stimuli were either broadband noise (BBN, containing complete binaural and spectral cues), low-pass filtered noise (LPN: <1 kHz, designed to contain only ITD information), band-pass filtered noise (BPN: 1/6 octave centred at 15 kHz, containing mainly ILD information), or high-pass filtered noise (HPN: >3 kHz, containing ILDs and spectral cues).

Across locations, ferrets were able to perform the task with each of the acoustic stimuli (binomial test against 50%, *p* < 0.001 for all ferrets and stimulus conditions, Table S1, Fig. 1C). Performance differences with limited cues were assessed with logistic regressions of stimulus type on trial outcome (with interactions) for each ferret (p < 0.05, Table S2). Performance in the band-pass condition (BPN, where localisation cues were restricted to narrowband ILDs) was significantly worse than the broadband condition (where all localisation cues are present) (Ferret F1302: across condition performance difference -11.6%, F1310: -7.5%, F1313: -6%, Fig. S1). Performance of ferrets in other limited cue conditions varied, two ferrets performed significantly worse in the low-pass condition when cues were limited to ITDs, although the magnitude of the difference was very small (F1302: -3.6%, F1310: -3.5%) and one ferret performed worse in the high-pass condition where there were no ITD cues (F1302: -6.8 %). We therefore predicted that any change in spatial tuning in A1 for BBN versus cue-restricted stimuli would be modest, and most marked for the BPN stimuli which consistently elicited worse performance across ferrets.

**Fig. 1:**
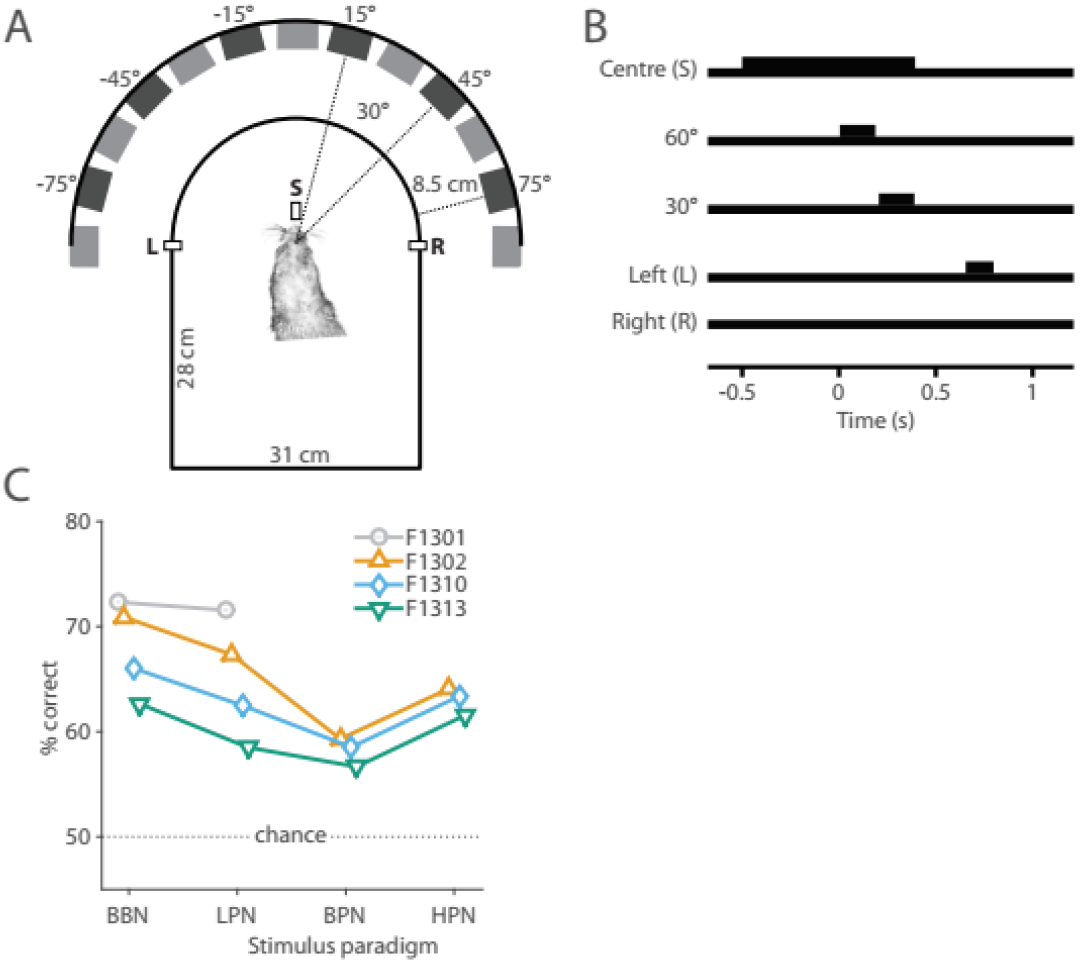
Ferret relative localisation behaviour. (A) Ferrets were trained to report the location of a target sound relative to a preceding reference, with target and reference sounds originating from two speakers separated by 30°. Ferrets reported the relative location of the target at a left (L) or right (R) response spout (positioned at ±90°). (B) Ferrets were required to maintain contact with the central start spout (S) for a hold time (500-1500 ms) before the reference was presented, and remain there until at least half way through the target stimulus before 170 responding. (C) Ferrets were tested in 5 different conditions in which the stimuli were broadband noise, (BBN) 171 low-pass filtered noise (<1 kHz, LPN), band-pass filtered noise (1/6 octave about 15 kHz, BPN) and high-pass 172 filtered noise (>2 kHz, HPN). All animals performed the task above chance in each condition (binomial test, p<0.001).

During task performance, we recorded from 398 sound-responsive units in A1 (total across all stimuli, ferrets, electrodes and recording depths, recording location confirmed with frequency tuning and post-mortem histology, Fig. 2C, D). Spatial tuning was calculated during task performance by considering the neural response to the reference sounds (Fig. 3A, B).

**Fig. 2:**
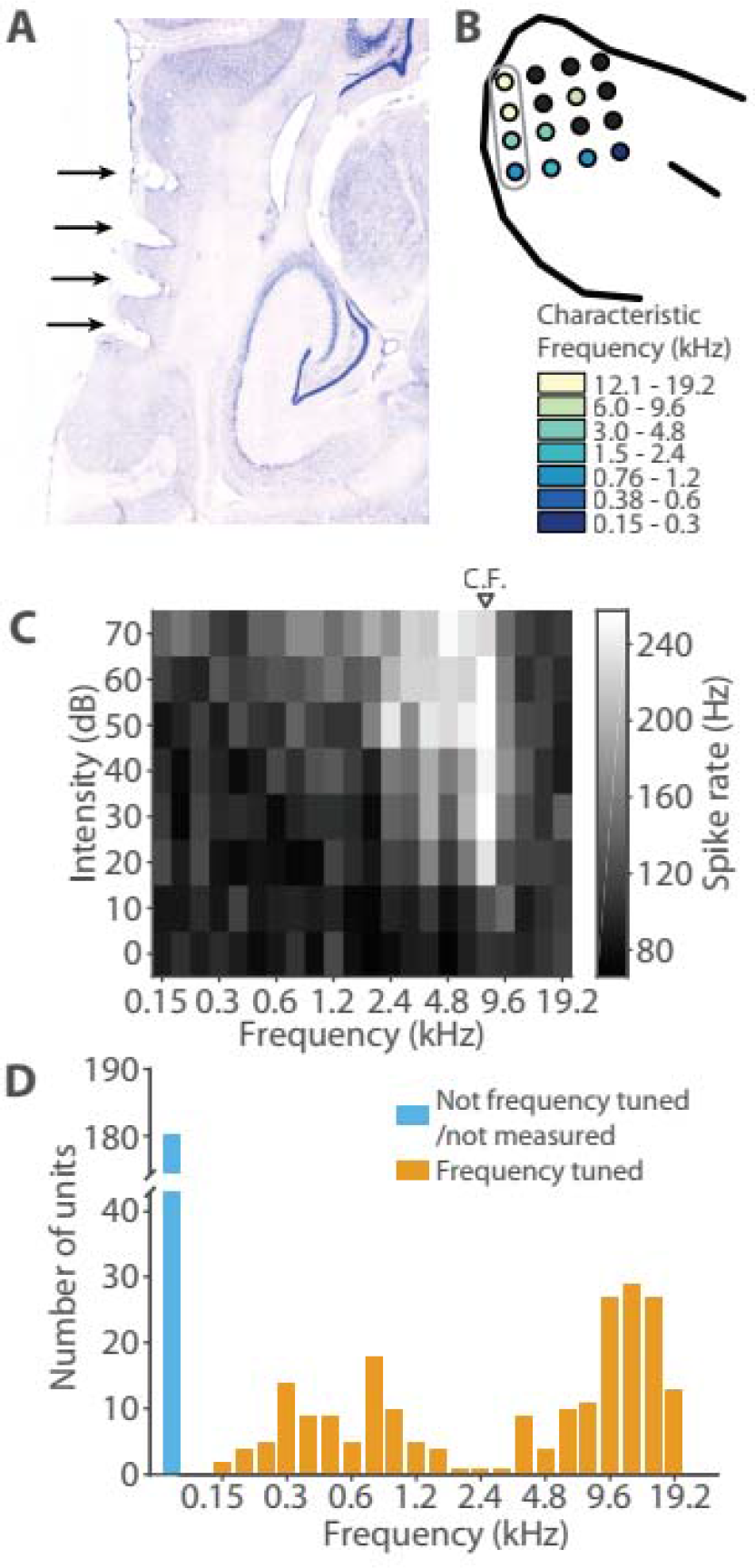
Experimental procedures. (A) Nissl-stained brain section of F1310 showing electrode tracks marked with electrolytic lesions (black arrows) of one column of the 4 x 4 electrode array. The locations of the electrode penetrations were clearly visible on the surface of the brain post mortem, indicated in (B) with each circle indicating the electrode sites. The circles are coloured indicating the frequency tuning of the units recorded from that electrode extracted from Frequency Response Areas (e.g. C), black filled circles indicates that no characteristic frequency could be estimated for that electrode. The grey outline indicates the electrode tracks shown in (A). D distribution of characteristic frequencies for all recorded units.

### Spatial Tuning in Auditory Cortex of Behaving Ferrets

To establish a benchmark for comparison, we first characterised the responses to broadband sounds containing a full complement of localisation cues. For each unit (e.g. Fig.3A, B), we constructed a spatial receptive field using the firing rate across stimulus presentation (0-150 ms; Fig. 3C). We defined units as spatially tuned if firing rates were significantly modified by location (Table 1, Kruskal-Wallis p<0.05). In total 253 units were responsive to BBN, of which 165 units (65%) were spatially tuned (Table 1).

**Table 1.** Spatial tuning properties of spatially modulated units.

|  | Sound responsive units | Spatially modulated units |  |  |  |  | Units with significant spatial information |  |
| --- | --- | --- | --- | --- | --- | --- | --- | --- |
|  |  | Number of units |  | Mean centroid (°) | Mean ERRF width (°) | Mean modulation depth |  |  |
| **BBN** | 253 | 165 | 65% | -27 ± 20 | 121 ± 12 | 39 ± 11% | 158 | 62% |
| **LPN** | 244 | 124 | 51% | -33 ± 21 | 119 ± 15 | 40 ± 15% | 109 | 45% |
| **BPN** | 97 | 50 | 52% | -25 ± 16 | 117 ± 12 | 41 ± 13% | 51 | 53% |
| **HPN** | 114 | 63 | 55% | -26 ± 23 | 122 ± 13 | 36 ± 10% | 48 | 42% |
| **CSS** | 128 | 94 | 73% | -33 ± 20 | 112 ± 16 | 50 ± 15% | 82 | 64% |

For each spatially tuned unit we described the preferred azimuthal direction by computing the centroid. The majority of units (147/165, 89%) had contralateral tuning (mean ± standard deviation (SD) left hem = 26.0 ± 18.2°, right hem. = −25.3 ± 24.1°, Fig. 3D). We also measured modulation depth and Equivalent Rectangular Receptive Field (ERRF) to determine tuning depth and width. Tuning was generally broad (Fig. 3E, mean ± SD ERRF width, left hem = 123.1 ± 10.0°, right hem = 119.5 ± 14.9°) with a diversity of modulation depths (Fig. 3F, mean ± SD modulation depth, left hem. = 38.1 ± 10.7%, right hem. = 39.9 ± 13.5 %). There were no significant differences in the distribution of centroid values (when considered relative to hemisphere of the unit), ERRF width or modulation depths between the left and right hemispheres (Unpaired T-test, p>0.05).

Spike timing conveys additional sound location information beyond that offered by spike rates (41–43). To test if our units conveyed information about sound location in the temporal pattern of spikes, we decoded spatial location using a pattern classifier based on Euclidean distances between firing patterns recorded in response to stimuli at each location, binned at 15, 50 or 150 ms resolutions (with 150 ms representing the firing rate across the whole stimulus). Classifier performance was summarised as the mutual information (MI) between actual and classified locations (Fig. 3G), and spatially informative units were identified as those with performance significantly greater than chance (permutation test, MI more than the mean + two standard deviations of MI calculated with shuffled sound locations, 250 iterations, p < 0.046). Using this measure, we found 62.5% (158 / 253) of units were spatially informative when decoded with at least one temporal resolution. Across units the best decoding window was equally distributed across bin widths (logistic regression of bin width vs. constant model, X^2^ = 2.11, p = 0.147, d.f. = 1, Fig. 3H).

While the best width of the decoder was equally distributed across units, the decoding performance (i.e. percentage of maximum available MI for perfect classification performance) was highest when we decoded sound location from responses binned in a single 150 ms intervals (i.e. a spike rate code, Fig. 3I). Statistical comparison confirmed a main effect of bin width (one-way ANOVA, F(2, 155) = 12.54, *p* < 0.001) with post-hoc contrasts (Tukey-Kramer corrected) confirming greater MI for decoding with 150 ms than 15 ms (*p* < 0.001) or 50 ms (p = 0.002) bins (Fig. S2).

**Fig. 3:**
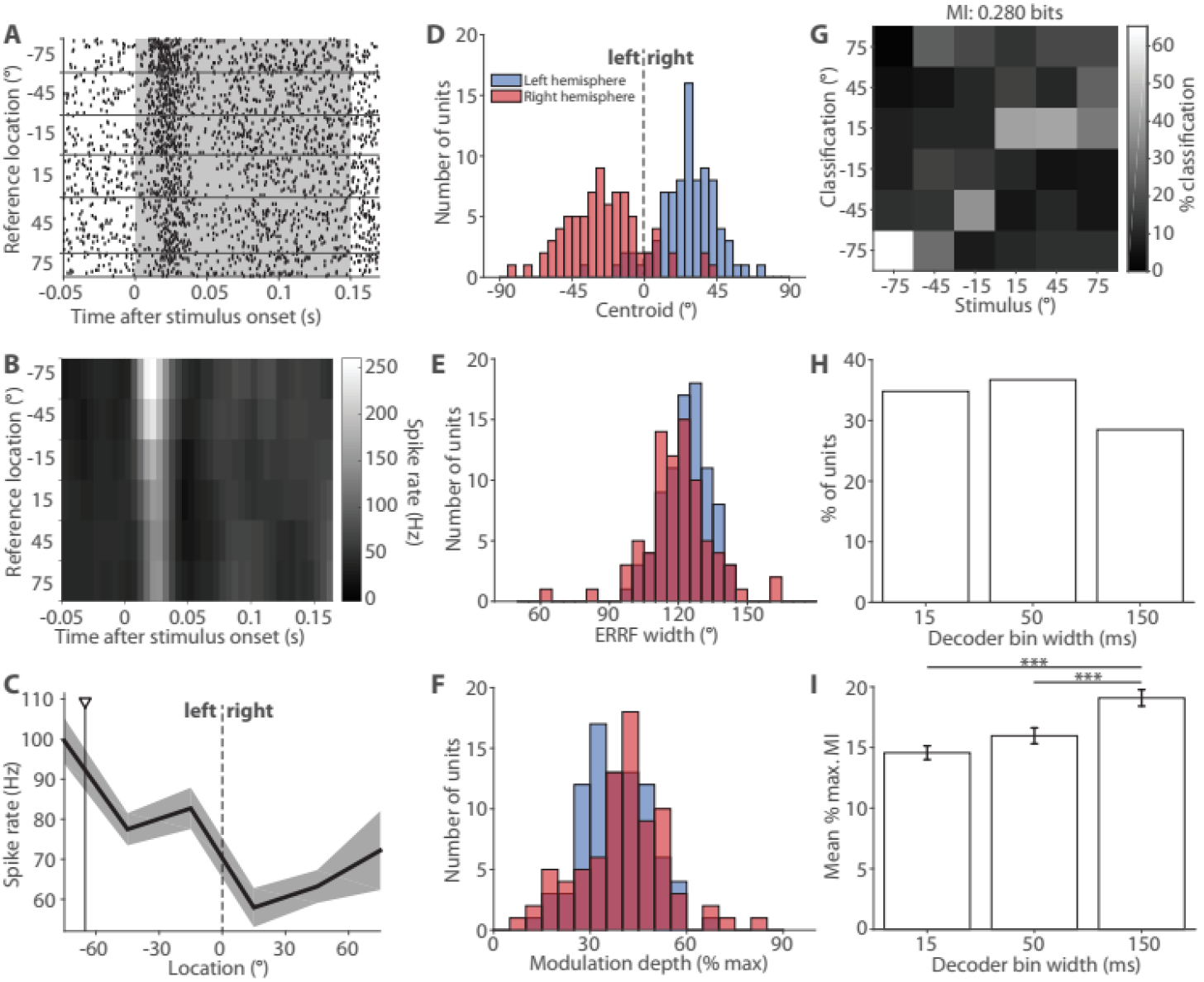
Units contain information about location of broadband noise bursts and are largely contralaterally tuned. (A) Example raster plot from one unit, with trials ordered by the location of the reference sound. The duration of stimulus presentation is indicated by the grey box. (B) Post-stimulus time histogram (spikes binned at 5 ms resolution) (C) Spatial receptive field (mean ± SEM spike rate calculated over 150 ms at each location. The unit shows clear contralateral tuning and a centroid of - 65° (black triangle/line). (D), Distribution of centroids (E) equivalent rectangular receptive fields (ERRF) widths and (F) modulation depths for all spatially modulated units (n = 165) split by left (blue, n = 85) and right (red, n = 80) hemisphere. (G) Stimulus location decoded with a Euclidean distance decoder on the spike response binned with 15 ms resolution for the unit in A-C. (H) Best decoder bin widths for all significantly informative units (158/253) (I) Mean maximum MI of significant units from (G) (n = 54, 58, 46). Statistics: (H) GLM: bin width. (I) one-way ANOVA Tukey-Kramer post-hoc pairwise comparisons. * p<0.05, ** p<0.01, *** p<0.001.

### Spatial receptive fields in A1 represent space, rather than localisation cues

We addressed the question of whether auditory cortex represents *space* or localisation cue values by contrasting the spatial tuning observed in response to broadband and cue-restricted sounds. We hypothesised that if A1 neurons represent auditory space their tuning should be independent of the spatial cue provided. In contrast, if A1 neurons encoded specific cues, spatial tuning should be abolished when the encoded cue was removed from the stimulus.

Individual units showed responses to cue-restricted stimuli similar to those observed with BBN stimuli (Fig. 4 A-C, Fig. S3 A). To assess the spatial tuning properties of all units, we compared the distributions of centroid, ERRF width and spatial modulation depth in responses to broadband, low-pass, band-pass or high-pass stimuli (Fig. 4 D-F, S3). We first compared the centroids of all units with spatially modulated responses to either broadband or limited cue stimuli. Centroids obtained for all cue limited conditions were no different to those measured with BBN (KS test p>0.05) nor between limited cue conditions (KS test, p>0.05), with the distribution of centroid differences being centred at zero (Fig. 4 D-F i, Fig. S3 B-D i). Tuning remained consistent when considering only the units that were spatially modulated in both conditions (paired T-test, p>0.05). Thus the direction of tuning was conserved across broadband, high-pass, band-pass and low-pass pass noise indicating that A1 neurons represented sound angle using both ILDs and ITDs in a redundant fashion.

Similarly for ERRF width, we found no significant differences when comparing receptive fields of units that were spatially modulated in any condition (T-test, p = 0.045, Bonferroni corrected alpha = 0.017), nor when comparing units that were spatially modulated in both conditions (paired T-test, p>0.05, Fig. 4E-F ii, S3 B-D ii). Finally, for units that were spatially modulated in any condition, there were significant differences in the modulation depth between responses to broadband and HPN (change in mean modulation depth: −4.7%, T-test, p = 0.004), but not for any other comparison (T-test, Bonferroni corrected p > 0.017). When comparing units that were spatially modulated in pairs of conditions, again there were no significant differences in between any conditions (paired T-test, p > 0.017, Fig. 4E-F iii, S3 B-D iii). Across the population of recorded neurons the changes in tuning observed were of the same order of magnitude to those observed when we compared repeated recordings of the same stimuli for the same units (Fig S4).

Thus removal of any localisation cue had no systematic effect on the spatial tuning properties of units in A1 that were spatially modulated in both BBN and cue limited conditions, suggesting that neurons can represent the location of stimuli using spatial cues in a redundant fashion, consistent with a representation of space rather than spatial cues.

**Fig. 4:**
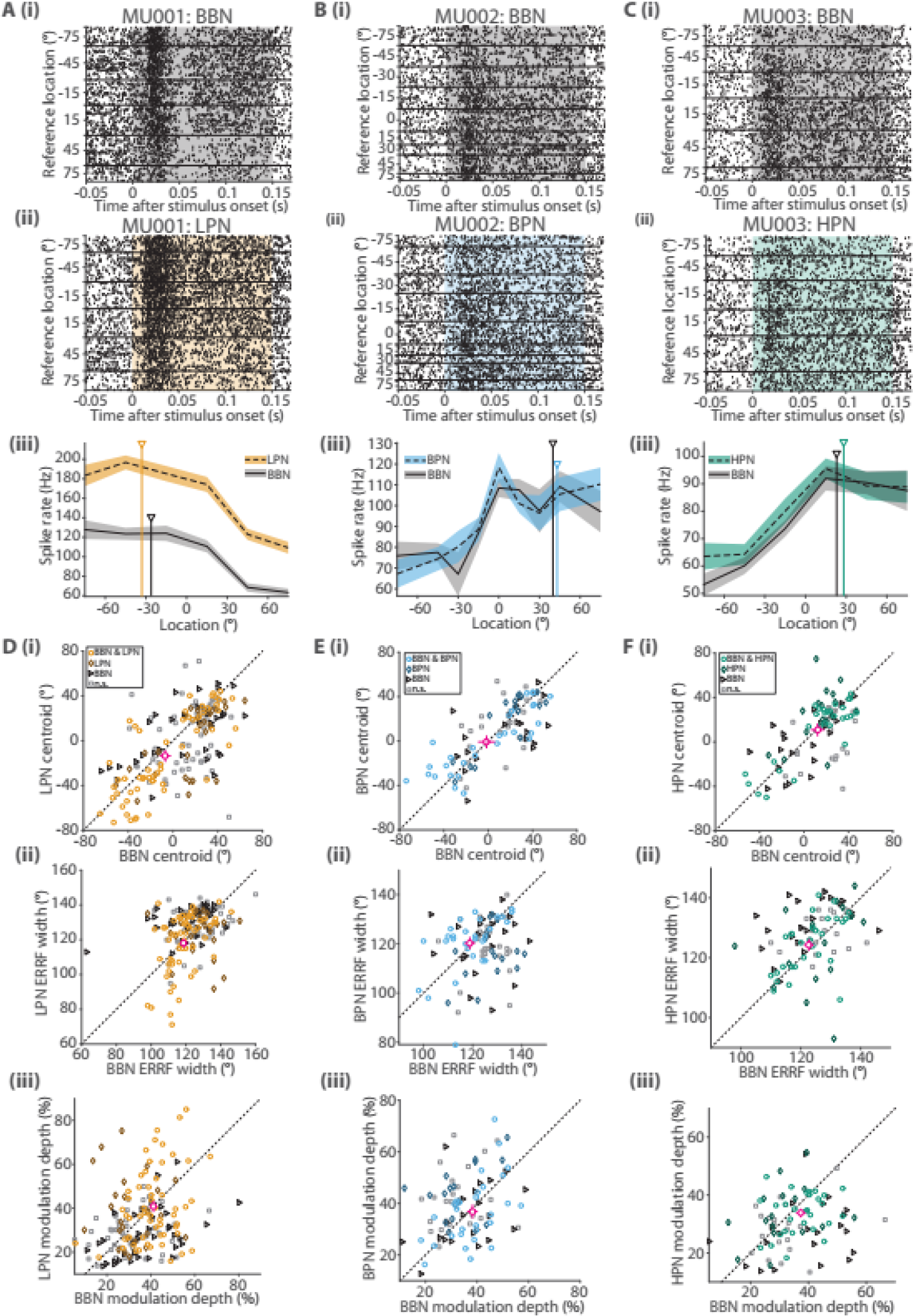
Changing spectral content of the stimuli had little effect on spatial tuning properties of units. (A)(i) Example raster from unit in response to broadband (BBN) stimuli. (ii) Raster of response to low-pass (LPN) stimuli of the same unit from (i). (iii) Spatial receptive field of the unit in response to BBN and LPN stimuli. (B, C) Same as (A) for BBN and band-pass (BPN, B) / high-pass (HPN, C) stimuli. (D) Centroids (i), ERRF width and modulation depth of units recorded in both BBN and LPN conditions. Units are marked according to whether they were spatially tuned in both conditions (circles), one condition alone (diamonds - LPN, triangles-BBN) or tuned in neither (squares). Mean ± standard error of the *jointly tuned units* (circles) is shown by crosshairs (circle, magenta). (E) Same as D for BBN and BPN stimuli. (F) Same as D for BBN and HPN stimuli.

### A subset of A1 units represent sound location across spatial cues

The above analysis suggests that spatial receptive fields are similar whether measured with BBN or limited cue stimuli. We next asked whether reducing the available localisation cues impacted the ability to accurately decode sound source location from neural responses, and whether spatial information was conveyed over similar timescales. Table 1 shows the proportion of units in which spatial location could be decoded significantly better than chance with each stimulus.

We first determined that cue-type did not impact upon the temporal resolution with which neurons best conveyed spatial information (logistic regression on unit classification (informative / uninformative), with bin width (15, 50 and 150 ms) and cue type (BBN, BPN etc.) as predictors revealed no significant effects: model vs. constant model, X^2^ = 6.25, p = 0.511, d.f. = 4, Fig. 5A).

**Fig. 5:**
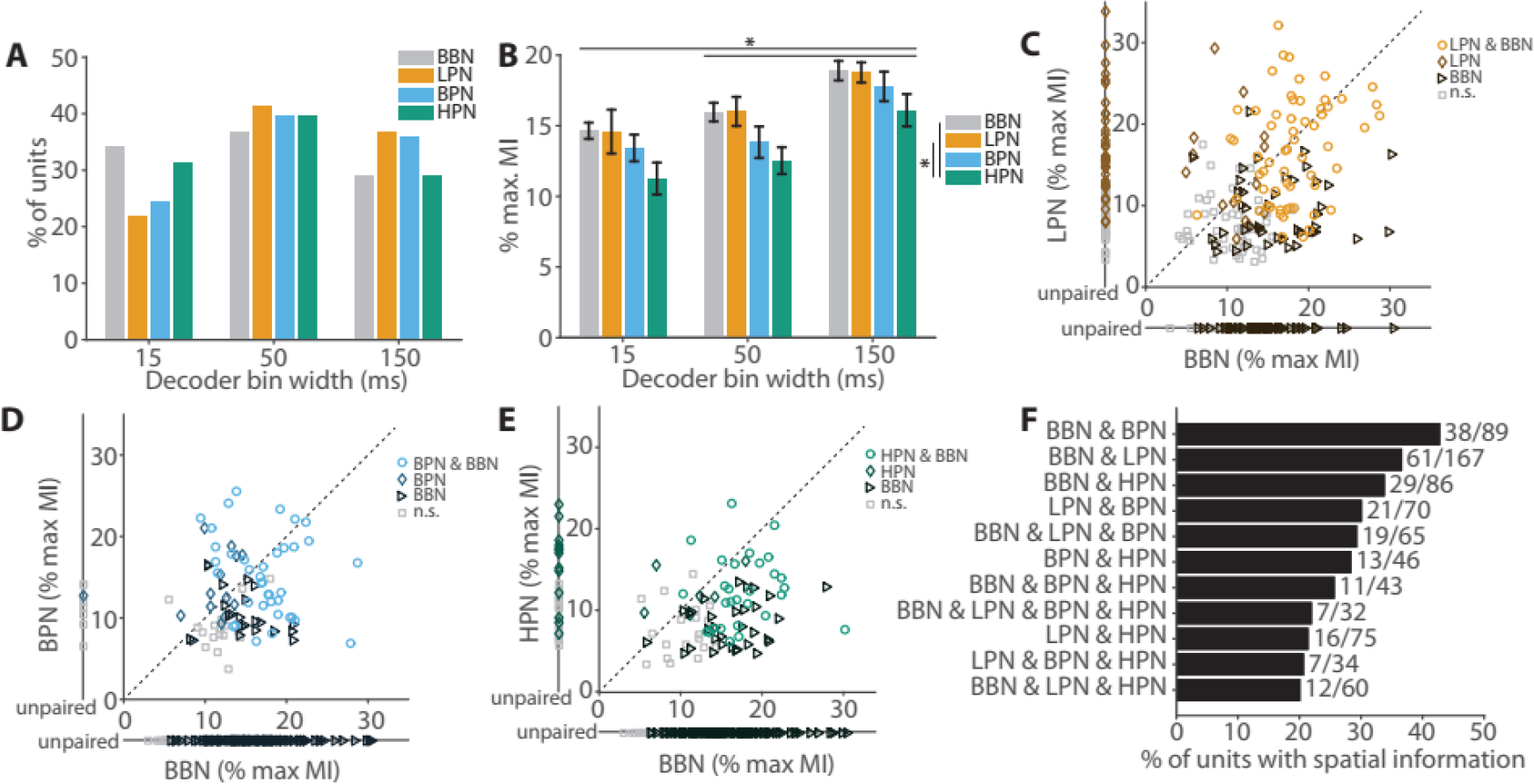
Comparison of spatial information in broadband vs. limited cue conditions. The amount of spatial information (MI) from stimuli with limited cues was compared with those where all cues were present (BBN). (A) Best decoder bin widths for all spatially informative units (BBN (n=158), LPN (109), BPN (51), HPN (48)). (B) Mean maximum MI of units from (A). (C) Comparison of MI in units recorded in both broadband (BBN) and (C) low-pass (LPN), (D) band-pass (BPN) and (E) high-pass (HPN) conditions. Units that were only recorded in one condition are plotted along a separate axis. (F) Proportion of units that showed information (in any bin width) in each combination of stimulus condition. Statistics: (B) Two-way ANOVA Tukey Kramer post-hoc pairwise comparisons.

We also compared best decoder performance (measured as the percentage of the maximum MI at the best bin width) across cue types (Fig. 5B). We found significant main effects of bin width (Two-way ANOVA, F(2, 354) = 21.57, *p* < 0.001) and stimulus condition (F(3,354) = 4.85, *p* = 0.003) but no interaction (F(6, 354) = 0.196, *p* = 0.978). Post-hoc tests (Tukey-Kramer, p<0.05) were consistent with the BBN data, with decoding being best when using a rate code. Additionally, decoding of responses to HPN was significantly worse than to BBN (Fig. 5E) and LPN conditions (Fig. S5B). To further investigate the changes in spatial information between stimulus conditions, we identified units that were spatially informative in both the BBN and cue-restricted conditions. Paired analyses revealed that for these units the amount of spatial information (at the best bin width) in BBN and LPN conditions, and in BBN and BPN conditions was equivalent (Fig. 5C & D, paired T test, *p* >0.05, N = 61 & 38 respectively). However, spatial information was significantly lower in high pass than broadband noise (Fig. 5E, paired T test, *p* < 0.001 (N = 29)).

To elucidate whether units were representing specific acoustic cues, or the spatial location of sounds independent of their underlying binaural cues we contrasted the number of units that were informative about sound location in more than one stimulus condition. Particular attention was paid to performance across conditions in which distinct binaural cues were presented (i.e. LPN, containing ITDs, and either HPN or BPN, which did not contain ITDs). We found that 30% (21/70) of units conveyed information about sound location across LPN and BPN and 21% (16/75) of units conveyed information across LPN and HPN ‒ conditions with mutually exclusive cue types. The ability of these units to represent sound location across these conditions argues that these units represent auditory space rather than acoustic cues and thus may be considered ‘cue-invariant’.

To further understand how the subset of ‘cue-invariant’ units represented sound location, we compared the spatial tuning properties of units that were spatially modulated in two conditions with mutually exclusive localisation cues (Fig. S3) and found that there were no significant differences in centroid for LPN vs BPN (ITDs vs ILDs, KS-test, p>0.05, N = 18) or for LPN vs HPN (ITDs vs ILDs & spectral cues, KS-test, p>0.05, N = 20). However, in units that were spatially informative in both LPN and BPN conditions, there was less information (−4.1%, measured as % max MI) in the BPN than LPN condition (Fig. S5, ILD vs ITD, paired T-test, p = 0.044, N = 21). For units that were informative in LPN and HPN conditions, there was less information in the HPN than LPN condition (−3.9%, ITD vs no ITDs, paired T-test, p = 0.035, N = 16) but there was no significant difference in information for units that were informative about BPN and HPN sound location (ILDs vs *no* ITDs, paired T-test, p = 0.627, N = 13). This is consistent with the ferrets being most impaired on BPN stimuli, and suggests that ITDs may provide more information about the location of a stimulus than ILDs or spectral cues.

In summary, many cells responded to multiple stimuli with mutually exclusive localisation cues suggesting that a proportion (25-30%) of spatially informative A1 cells can generalize the location of sound sources across their component acoustic cues.

### A competing sound source sharpens spatial receptive fields but does not affect spatial information content

If neurons encode auditory objects rather than simply auditory space then competition between objects might be expected to refine spatial tuning (28, 32–34). We hypothesized that a competing sound source would increase spatial sensitivity of neurons in auditory cortex by causing competition between two auditory objects. To test this, we repeated recordings in the presence of a competing sound source (CSS), a broadband noise presented at 0° azimuth and +90° elevation whose level was randomly stepped within a range of ± 1.5 dB SPL every 15 ms to aid segregation from the target stimuli.

Adding a competing source resulted in a mild impairment in relative localization in one of the three animals tested (F1310: mean change: −5.8%, Fig. 6A, GLM on response outcome with stimulus as predictor, p = 0.011, Table S2). At the neural level, there was no effect of the competing source on the direction of spatial tuning in units that were spatially modulated in both conditions (Fig. 6E, KS-test, p = 0.580). However, adding the competing source did sharpen spatial tuning, as ERRFs widths were narrower (paired T-test, p = 0.002) and modulation depths greater (p < 0.001) when the competing source was present (Fig. 6F, G). These results are consistent with auditory cortex encoding the location of auditory objects since spatial tuning was enhanced (as assessed by enhanced modulation depth and smaller tuning width).

We also assessed how adding a competing sound source affected decoding of spatial information from spiking responses. We found that adding the competing source did not affect the proportion of units with best decoding performance in each bin width (Fig. 6H, X^2^ = 0.0765, p = 0.4812, d.f. = 2). Adding the competing source did not change the information content of units in each bin width (Fig. 6I-J, two-way ANOVA, effect of CSS: F(1, 234) = 0.78, *p* = 0.378, Fig. 6I), nor was there an interaction between bin size and CSS (F(2, 234) = 0.98, *p* = 0.376). There was a main effect of bin width (F(2, 234) = 13.48, *p* < 0.001) reflecting the greater information for 150 than 15 or 50 ms bin widths (Tukey-Kramer, p<0.05).

**Fig. 6:**
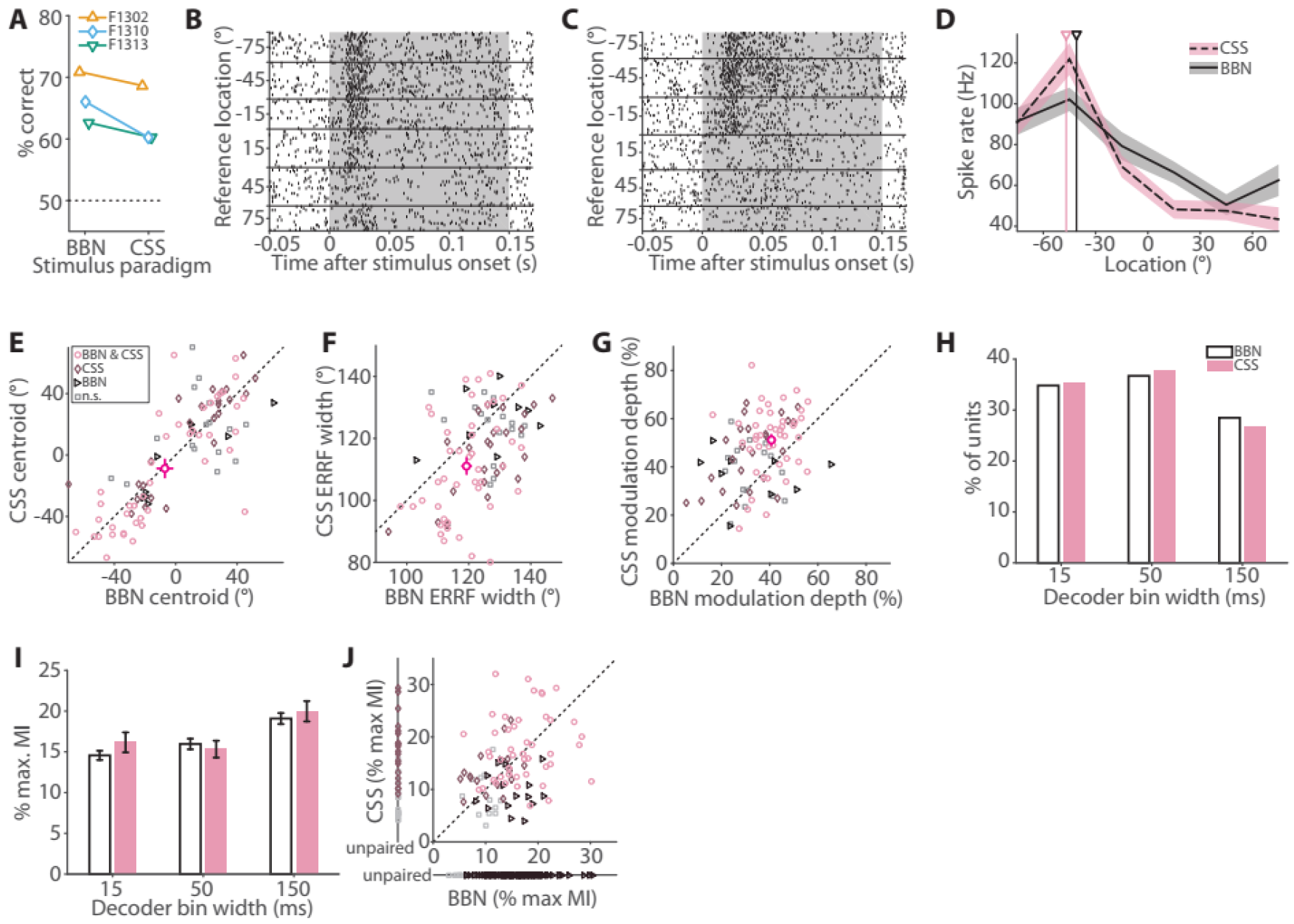
A competing sound source affects spatial tuning properties but has no effect on spatial information content. (A) Performance in the relative localization task in broadband conditions (BBN) and in the presence of a competing sound source (CSS) for each ferret. (B) Example raster in response to BBN stimuli and (C) CSS stimuli. (D) Spatial receptive fields of the same unit with centroids indicated by triangles (E) Centroids, (F) ERRF widths and (G) modulation depth of units recorded in both BBN and CSS conditions. Mean ± standard error of the mean of each is shown by crosshairs. (H) Distribution of best bin widths for all significantly spatially informative units (BBN n = 158, CSS n = 82). (I) Mean maximum MI of units from (G). (J) Comparison of MI in units recorded in BBN and CSS conditions. Legend as E.

### Encoding of auditory space by populations of neurons resembles a labelled-line model

While individual units conveyed significant spatial information, they still fall some way short of perfect performance (mean decoding performance only reached between 15 and 19% of the maximum MI possible) implying some sort of population coding is necessary to reconstruct sound location perfectly. To address this we used two models of population decoding to reconstruct sound location from neural activity (Fig. 7A): (1) A labelled-line model that decoded sound location from the activity pattern of neurons with heterogeneous spatial tuning, and (2) a two-channel model that compared the summed activity of neurons in each hemisphere of the brain (hemispheric channel) or summed activity of two populations of neurons with centroids in left and right space respectively (opponent channel). These models reflect the predominant theoretical descriptions of how neural circuits compute sound location (17, 36–39) and we can gain further insight into how sound location is represented in A1 by determining the success of these models in decoding sound location.

To assess whether the spatial receptive fields of units we recorded from A1 were more consistent with a labelled line or two-channel code, we first compared predicted and observed distributions of correlation coefficients (R) calculated between tuning curves obtained for all pairs of units. The two-channel models both predicted distinct peaks in distributions at R = ±1, as tuning curves should be strongly correlated within (positive correlation) and between (negative correlation) channels (Fig S6B). In contrast, the labelled-line model predicted a graded distribution of correlation coefficients, with many unit-pairs falling between -1 and 1 (Fig S6A) (17). We found that the distribution of correlation coefficients for units recorded in A1 most closely resembled the labelled line model with correlation coefficients distributed broadly between +1 and -1 (Fig S6C-G).

We next used maximum likelihood decoders (similar to ref. 39) to estimate sound location on single trials using the joint distributions of spike rates (a) recorded across units from each hemisphere (hemispheric two-channel), (b) recorded across units tuned to each hemifield (opponent two-channel), or (c) of individual units (labelled-line). The labelled-line decoder substantially out-performed the hemispheric and opponent two-channel decoder in all stimulus conditions (Fig. 7B-E) with little difference between stimulus conditions. The labelled line decoder reached > 85% with a minimum of 20 units (CSS condition) and maximum of 40 units (HPN) whereas neither two channel model exceeded 45% correct. There was little difference between the performance of the opponent and hemispheric two-channel models, possibly because the majority of neurons were tuned contralaterally (89% of units in response to BBN).

**Fig. 7:**
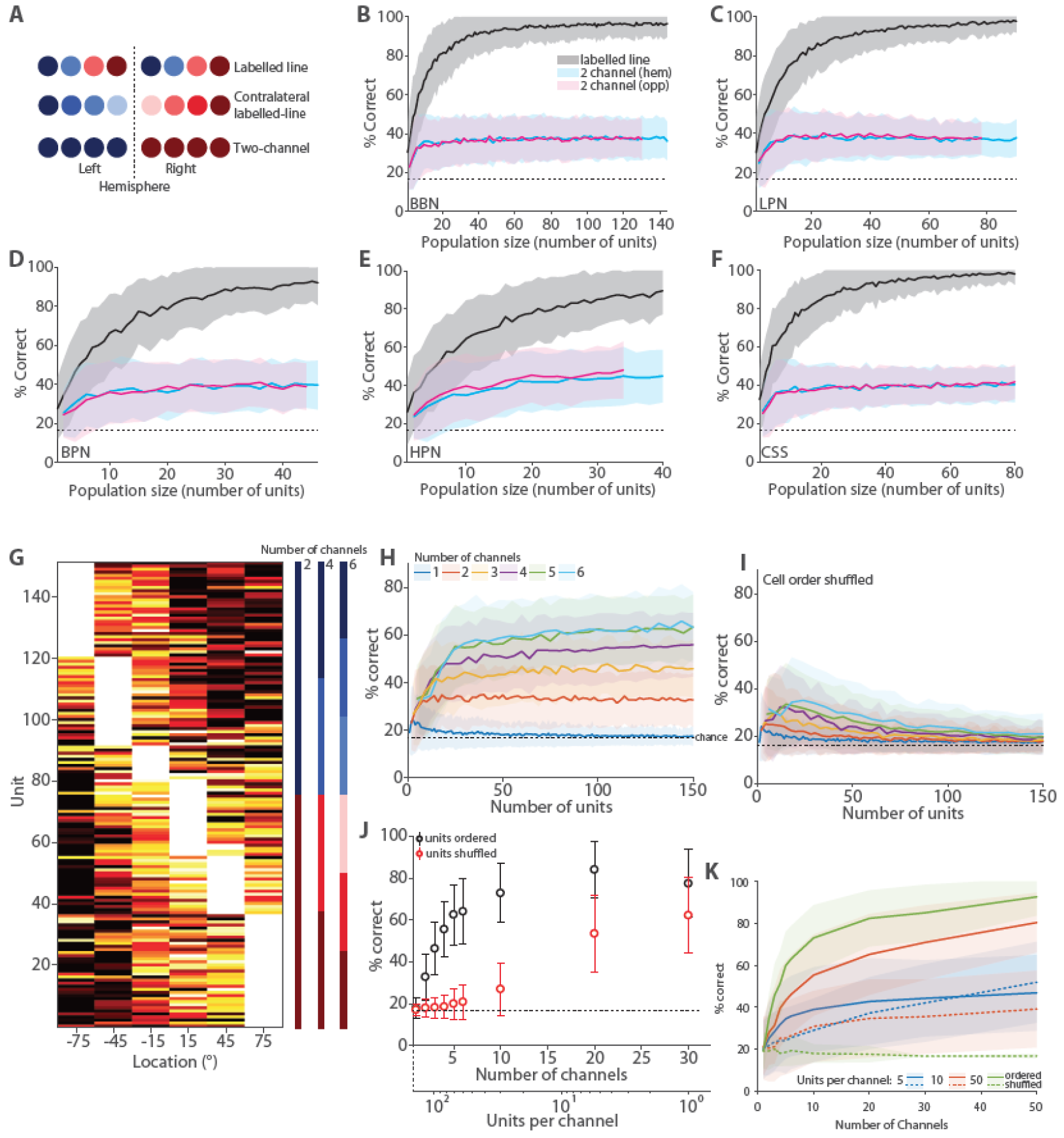
Population labelled-line decoding outperforms two-channel models. A schematic of three possible spatial coding mechanisms B-F, Performance of the two-channel (hemispheric—blue, opponent - magenta) and labelled line (grey) decoders in each stimulus condition from populations of neurons that had significant MI about location. Performance of each decoder was calculated as percentage of correctly classified individual trials. The dotted line indicates chance (1/6). The black line and coloured bands indicate the mean ±SD of the performance for 250 different random sub populations of each size drawn from the full sample of cells. G spatial receptive fields for all units used in the population analysis, ordered by best position. H decoder performance for models with increasing channel numbers with units grouped according to G, I decoder performance when units are randomly ordered and assigned to channels, J, comparison of decoder performance and shuffled performance for a fixed population of 140 cells, and increasing channel number. K, Effect of the number of units per channel using data from modelled unit responses for ordered and shuffled units (see methods)

Previous work has suggested that spatial representations in auditory cortex may depend on more than two channels. For example, it has been posited that there might be a third central channel (44, 45) while others have suggested many channels with 6° widths tiling space (46). In order to investigate this idea further, we modified our two-channel decoder to compare decoding performance of neural populations divided into *N* channels according to spatial tuning observed in response to BBN stimuli. Cells were ordered by the location of the peak firing rate and divided into channels with equal numbers of units per channel (Fig. 7G). As the number of channels increased, decoding performance also increased (Fig. 7H). If the order of the units was randomly shuffled so that units within a channel were sampled without regard to their spatial tuning, performance was at chance levels, with the exception of populations containing very few units per channel (Fig. 7I).

When population size was held constant and the number of channels increased (Fig. 7J, for a population of 140 units), shuffled performance was always worse than ordered performance. However, as the number of channels increased, the number of units per channel decreased, and thus the effect of shuffling unit order diminished. In the extreme case, where each channel had one unit, the shuffled and ordered distribution differed only in the relationship between channels and thus both shuffled and ordered populations provided labelled line systems. In order to further understand the relationship between the number of channels and the number of units per channel, we simulated responses of cells based on the spatial receptive fields of units that responded to BBN stimuli (Units in Fig 7J). This allowed us investigate the effect of increasing the number of channels while keeping the number of units per channel constant (Fig 7K).

For channels containing similarly tuned units (i.e. ordered) increasing the number of channels rapidly improved decoding performance up to ~20 channels, after which performance saturated. For high channel counts (e.g. n > 30) and low numbers of units per channel (n = 5), decoding performance of shuffled and ordered populations converged. This replicated our findings for the labelled line decoder using recorded neurons (Fig. 7J). However when the number of units per channel was high (n = 50), decoding was far superior when units were ordered than when shuffled.

These results can be explained by a dependence of the labelled line decoder on heterogeneity in the underlying tuning functions of individual cells and channels. When the number of units per channel was small and the number of channels was large, the diversity of spatial tuning curves present across units was preserved across spatial channels and therefore the decoder remains successful in reconstructing sound location. This is true regardless of the order of units because, with few units per channel, there is little averaging across differently tuned units within channels in the shuffled analysis. In contrast, when there are many *units per channel*, the order in which units are arranged before integration into channels is critical. When units are ordered by spatial tuning, adjacent units have similar spatial receptive fields and therefore the effect of integration is primarily to decrease noise, while channels remain strongly tuned to sound location despite averaging. Thus the heterogeneity of individual units is preserved at the channel level, while decoding performance improves due to reduction in noise. In contrast, when units are shuffled, adjacent units have differing spatial receptive fields and so the effect of integration is to average out spatial tuning. Channels in the shuffled decoder are therefore weakly tuned relative to the underlying units and decoding performance is poor.

## Discussion

We sought to understand how available localisation cues affect encoding of azimuthal auditory space in primary auditory cortex of ferrets performing a relative sound localisation task. Reducing the available localisation cues (ITD, ILD and spectral) had a weak effect on the animals’ behaviour, and on the spatial tuning of neural units in A1. A subpopulation of units were capable of representing auditory space in a cue-invariant manner, while the behaviour of other units was more consistent with the encoding of a specific localisation cue. We found that tuning curves of units in each hemisphere had best azimuths distributed across the contralateral hemifield, and were narrower than would be expected for a two-channel model of spatial representation, consistent with a labelled-line model of sound location encoding. Labelled-line and two-channel decoders based on the spike rates of populations of neurons both performed above chance in all stimulus conditions but the labelled-line decoder greatly outperformed the two-channel decoder.

Ferrets were able to perform the relative localization task using both ILD and ITD cues when tested with high pass, band-pass and low-pass noise. In such conditions, directional preference of cells (as measured by centroids) was stable and we identified a subset of roughly a quarter of units that were significantly spatially informative when provided with either ILDs or ITDs. This suggests a subpopulation of A1 cells provide a cue-invariant representation of sound location. Other units only conveyed spatial information in the presence of ITDs or ILDs. Similar results have been obtained using human neuroimaging where regions of cue-independent and cue-specific voxels were observed (26). These neuroimaging results suggest that a representation of azimuthal space exists within auditory cortex that is independent of its underlying acoustic cues. Our results provide the first cellular resolution evidence in support of this hypothesis in cortex of behaving subjects. It will now be important to ask how binaural cues are integrated within A1, and, given recent observations about world-centred spatial tuning in A1 (11), what reference frame cue-invariant neurons operate in. Since many spatially tuned units were informative about sound location only when specific cues (ILDs or ITDs) were present, it is possible that the responses of cue-specific units are integrated within A1 to develop cue-tolerant responses.

The addition of a competing sound source sharpened spatial receptive fields by narrowing and increasing the modulation depth consistent with previous findings in songbirds (33). Despite these improvements in representation of spatial location, there was no overall difference in the amount of spatial information conveyed by units in the presence of a competing sound source, although population responses saturated with a smaller number of units than for other conditions suggesting narrower tuning improves the resolution with which space can be decoded. It may be that the observed changes in tuning enable the animal to selectively attend to the behaviourally relevant source while ignoring the competing sound. If true, we would predict that blocking these adaptive changes in spatial tuning would lead to large deficits in relative sound localization. Such experiments will require more detailed knowledge of the circuit mechanism through which spatial tuning in the auditory system is modulated.

We observed that a labelled line decoder outperformed a two-channel decoder, and that decoding ability scaled with the number of spatial channels. Taken together with the observations that (a) centroids tiled auditory space and (b) spatial tuning curves were narrower and less correlated across units than a two-channel model would predict, these data support the labelled line model of spatial encoding. However, rather than each hemisphere containing a full representation of auditory space (as found in gerbil, 39) each hemifield of space was represented by a labelled line system within the contralateral A1. This is consistent with demonstrations of contralateral tuning bias in anaesthetised, passively listening or awake behaving cats (37,47,12), awake monkeys (10) and anesthetised ferrets (43). Here we build on these findings by suggesting that within-hemisphere tuning supports multiple spatial channels - rather than two channels each representing left and right space as suggested elsewhere (37). A contralateral labelled line code can explain why unilateral inactivation of A1 selectively impairs localisation in the contralateral hemifield (13, 14, 40). Several questions arise and must be answered in the future: How does midline tuning emerge in non-primary fields (48) to enable the brain to locate or track sounds crossing the midline, and what are the effects of unilateral hearing loss on cortical representations of sound location (as distinct from localization cues)?

We tested a modified version of the labelled line code in which increasing population sizes were tested for models with increasing numbers of channels. This design allowed us to address the possibility that there may be more than two channels (44–46), and assess the impact of within-channel averaging on model performance. Thus we made a version of the labelled line decoder that compared channels of small populations of similarly tuned units that were summed together. We found that as the number of channels increased, decoder performance increased, lending further support to a labelled-line type encoding of auditory space in A1. Importantly, shuffling the spatial tuning of units decreased decoder performance in all cases, although above chance performance was observed where heterogeneity between the channels was maintained by having very small numbers of units per channel. This was consistent with the idea that averaging heterogeneous spatial receptive fields leads to loss of information (49, 50).

A labelled line encoding of auditory space is consistent with the formation of auditory objects in A1, and would allow distinct subpopulations of neurons to represent each object (34). This would require combination of the separate spatial cues that derive from the same object to form a channel-based representation of auditory location as observed behaviourally in humans (46). This process might begin in the midbrain where there is evidence of integration of spectral cues and binaural cues (24, 25, 51). Determining whether auditory cortex forms representations of individual discriminable auditory objects (or sources) will involve testing under more naturalistic listening scenarios where there is variation across multiple orthogonal properties (26, 52, 53) and the use of multiple sound sources in the context of a behavioural task.

## Acknowledgments

This work was supported by a Wellcome Trust and Royal Society Fellowship (098418/Z/12/Z) and a BBSRC project grant BB/H016813/1 to JKB and a UCL Grand Challenges Studentship to KCW.

## Methods

### Animals

All animal procedures were approved by the local ethical review committee and performed under license from the UK Home Office in accordance with the Animal (Scientific Procedures) Act 1986. Four adult, female, pigmented ferrets (*Mustela putorius furo*) were used in this study, 1-3 years old. All animals received regular otoscopic examinations to ensure that both ears were clean and healthy.

### Behavioural Training and Testing

The ferrets were trained to perform a relative localisation task similar to that performed by humans in (54). Ferrets were trained to report the location of a target sound relative to a preceding reference sound presented from a location either 30° to the left or right of the target (Fig. 1B). Training stimuli were broadband noise with the sound levels on each trial roved (55, 58 or 61 dB). To maximise the number of trials per location for neural recordings, once trained and implanted ferrets were tested at a single sound level. An animal was considered ‘trained’ when they reached criterion performance of 65% correct on the training stimuli (chance performance in this two-alternative task is 50%). Once trained, ferrets were chronically implanted with electrode arrays and subsequently tested with cue-restricted sounds and with competing sound sources.

For cue-restricted testing subjects were presented with broadband noise at a single sound level (BBN), low-pass noise (LPN) where the stimuli were low-pass filtered (finite-duration impulse response (FIR) filter with cut-off 1 kHz with 70 dB attenuation at 1.2 kHz), band pass noise (BPN, filter width one sixth octave around 15 kHz; 47 dB attenuation at 10 kHz and 55 dB attenuation at 20 kHz), high-pass noise (HPN filter cut-off 3 kHz; 70 dB attenuation at 2 kHz), All stimuli were presented at 61 dB SPL (measured at the position of the ferret’s head). The low-pass stimuli were designed to present the ferret with ITD cues only, the band-pass stimuli to provide ILD cues with very limited spectral cues, at this narrowband frequency ferrets rely on ILD cues to localise sounds (Keating et al., 2014) and the high-pass stimuli to exclude ITD cues but maintain ILDs and spectral cues.

BBN stimuli were also presented with a competing sound source (CSS) which consisted of continuous broadband noise whose amplitude was stepped (randomly drawn from a Gaussian distribution of values with mean 55 dB SPL) every 15 ms presented from a single speaker located directly above the animal when its head was at the start spout. Target stimuli were presented at varying levels (55, 58, 61 dB SPL).

Ferrets were tested twice daily Monday-Friday with each stimulus condition being tested in two sessions on the same day over a two week period. At the beginning of each week, ferrets were run on the training stimuli and only proceeded to testing when they reached criterion (65%).

### Electrode implants

Arrays of 16 independently moveable tungsten electrodes in a WARP-16 device (Neuralynx Inc., Bozeman, MT), were implanted into left and right auditory cortex during an asceptic surgery. Ferrets were provided with appropriate pre, peri and post-operative analgesia under veterinary advice. After surgery electrodes were moved into the brain and subsequently by 50-150 μm whenever the ferret had completed a full cycle of behavioural testing. In this manner over the course of 1-2 years recordings were made from each cortical layer in each ferret. Testing was complete when the electrodes had moved a depth that exceeded the estimate for the maximal depth of auditory cortex (2 mm). The location and final depths of the electrodes were later confirmed by histology. These data, combined with estimates of frequency tuning made at each site, enabled an estimate of the location of each electrode in auditory cortex (Fig. 1).

### Neuronal Recording

Signals were recorded, amplified up to 20,000 times and digitised at 25 kHz (PZ5, TDT). Data acquisition was performed using TDT System 3 hardware (RZ2), together with desktop computers running OpenProject software (TDT) and custom scripts written in MATLAB. Headstage cables were secured to custom made posts which screwed onto the protective caps that housed the implants, allowing the ferret free movement within the chamber.

### Data analysis

#### Spatial tuning features

Clusters identified in the spike sorting (see Supplemental methods) were deemed sound responsive if the spike rate 50 ms post-stimulus onset was significantly different than the baseline firing rate in the 50 ms preceding stimulus onset (paired t-test, p<0.05).

Spatial receptive fields were defined by calculating the mean spike count relative to baseline (mean response 150 ms prior to stimulus onset) of the response over stimulus presentation at each location of the reference sound (150 ms). A unit was defined as spatially tuned if its response was significantly modulated by location (Kruskal-Wallis test, *p* < 0.05).

The best azimuth (peak of the spatial receptive field) of each unit was given by its centroid and the location of the peak firing rate. The centroid was calculated in a similar way to Middlebrooks and Bremen (34): Firstly the peak rate range was determined as one or more contiguous stimulus locations that elicited spike rates within 75% of the unit’s maximum rate plus one location on either side of that range. The locations within the peak range were treated as vectors weighted by their corresponding spike rates. A vector sum was performed, and the direction of the resultant vector was taken as the centroid. The best azimuth was defined at the location of the peak of the spatial receptive field.

The breadth of spatial tuning of a unit was represented by the width of its equivalent rectangular receptive field (ERRF, Lee and Middlebrooks, 2011), which corresponds to the width of a rectangle with a total area the same as the area under the spatial receptive field and height equal to the peak firing rate. Modulation depth was defined as ((max response ‒ min response) / max response) × 100.

Differences between the centroids in each stimulus condition were tested with a two-sample Kolmogorov-Smirnov test (p<0.05). ERRF widths and modulation depths in each of the stimulus conditions tested with paired or unpaired T-tests (p<0.05) depending whether the comparison was across all units or between the same units respectively.

#### Spike pattern decoding of individual units

Responses were binned with 15 ms, 50 ms or 150 ms resolution across the duration of the reference sound presentation. Spike patterns were decoded by comparing a single test trial PSTHs to template PSTHs calculated as the mean PSTH for each location being tested (with the to-be-tested trial excluded (55). Test trials were classified by location according to the lowest Euclidean distance between test PSTH and template PSTHs. Mutual information (MI) was calculated between the actual and decoded sound location to quantify decoder performance. To test for significance, bootstrap simulations (250 repeats with resampling) on shuffled data were performed. The MI was deemed significant if it was higher than the mean of the shuffled distribution plus 2 standard deviations. Since in some stimuli (BBN and BPN) extra locations were tested (see stimuli and speaker array), the percentage of the maximum MI possible (log2 of the number of locations tested) was calculated in order to compare decoder performance across conditions with different speaker numbers.

### Population decoding

Similar to Belliveau et al. (39), a Bayesian maximum likelihood decoder was implemented to test different models of location coding. Bayes rule states that the probability of a stimulus given firing rate *x* is:

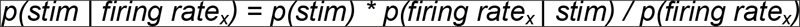

Two models were tested for decoding sound location: The labelled-line and the two-channel hemispheric model. For the labelled-line model, the probability of a given firing rate in each neuron in the population (N) given stimulus *Y* was calculated as the product of the probabilities of the firing rate of each cell (i):

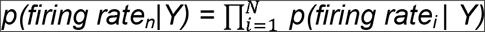

For the two-channel hemispheric model, two populations of neurons were defined by the hemisphere of the brain they were recorded from, for the opponent-coding model, two populations of neurons were defined by the side of space their centroid occupied. The likelihood term was calculated using the mean firing rate across all neurons in the population at each location. The joint probability of a set of firing rates in each hemisphere was calculated as the product of the probability of the mean firing in each hemisphere:

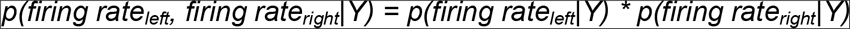

For the modified labelled line model, the whole population of neurons ordered by best azimuth was divided into channels (1-30 channels) with equal numbers of units. The likelihood term was calculated using the mean firing rate of all neurons in each channel at each location. The joint probability of a set of firing rates in each channel (i) was calculated as the product of the mean firing in each channel (for N channels):

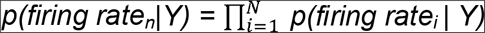

Since the number of presentations of a stimulus in a given recording sessions was not equal, the probability of the firing rate of each neuron was calculated using the maximum number of equal trials (randomly selected without replacement) for each location so that for the purposes of the population decoder, each location was presented with equal probability thus *p*(stim | firing rate_Y_) ∝ *p*(firing rate_Y_ | stim) therefore, the likeliest source location was defined as the max *p*(firing rate | stim). The models were tested for populations increasing in size from 1 to the maximum number of units in each stimulus condition with units randomly drawn from all the available units with replacement (bootstrap resampling). For each population size, a single trial from each cell was selected for decoding. This process was repeated 250 times for each population size. For each unit of the population, the mean and standard deviation of the spike counts from trials at each azimuth was calculated. Any unit recordings with fewer than 7 trials at any location were excluded from the population testing. Units were selected from recordings where the locations tested spanned -75° to 75° in 30° steps and where at least one of the bin widths had significant MI in the spatial location decoding. Where multiple test sessions were made at the same recording site, the best recording was defined as the unit with the highest MI value.

For testing the number of channels with a fixed number of units (Fig 7L), linear interpolation between the nearest actual values was performed so that the same, maximum number of units could be compared. For testing the effect of channel number while the number of units per channel was maintained (Fig 7K), we fitted Gaussian tuning curves to the population of units that responded to BBN and fulfilled the above criteria. For each population size (i.e. the number of channels and number of units per channel necessary), Gaussian tuning curves were selected from the population (with replacement) and 100 trials at each location were generated by drawing spike rates from Gaussian distributions with mean and variance of the fitted curves at each location. These data were subsequently used in the same way as data recorded from each cell in the modified labelled-line decoder.

## Supplementary Methods

### Behavioural Task

The ferrets were trained to perform a relative localisation task similar to that performed by humans in (54). Ferrets were trained to report the location of a target sound relative to a preceding reference sound presented from a location either 30° to the left or right of the target (Fig. 1B). The behavioural task was positively conditioned using water as a reward. During testing days ferrets did not receive any water in their home cage, although where necessary the water obtained during testing was supplemented to a daily minimum with wet food.

On each trial, ferrets nose poked at a central port to initiate stimulus presentation after a variable delay (500 – 1500 ms). Trial availability was indicated by a flashing LED (3Hz) mounted outside the chamber, behind the plastic mesh that enclosed the chamber, approximately 15 cm from the floor of the chamber and flashed. Following sound presentation, subjects indicated their decision by responding at a left or right reward spout. Responses were deemed correct the animal responded left when the target was to the left of the reference, and right when the target was to the right of the reference. Ferrets always received a water reward for correct responses from the response spout and received an additional reward from the start spout 5% of the time. Incorrect trials resulted in a 7 s time out before the next trial could be initiated, and were followed by correction trials (which were excluded from analysis) in which the stimulus was repeated. All training and testing was fully automated with all sensor input, stimulus presentation and solenoids coordinated via TDT System III hardware (RX8; Tucker-Davis Technologies, Alachua, FL) and custom-written circuit running in Open Project (TDT Software) and MATLAB (MathWorks Inc., Natick, USA).

### Stimuli and Speaker Array

All stimuli were noise bursts generated afresh on each trial in Matlab at a sampling frequency of 48 kHz. Sound stimuli were presented from thirteen loud speakers (Visaton SC 5.9) positioned in a semicircle of 24 cm radius around one end of the testing chamber (Fig. 1A), speakers were evenly positioned from -90° to 90° at 15° intervals approximately at the height of the ferret’s head when at the central start spout. Speakers were calibrated to produce a flat response from 200 Hz to 25 kHz when measured in an anechoic environment using a microphone (Brüel and Kjær 4191 condenser microphone). The microphone signal was passed to a TDT System 3 RX8 signal processor via a Brüel and Kjær 3110–003 measuring amplifier. Golay codes were presented through the speakers and the spectrum was analysed and an inverse filter was constructed to flatten the spectrum (Zhou, 1992). All sounds were presented low-pass filtered below 22 kHz (FIR filter <22 kHz, 70 dB attenuation at 22.2 kHz) and with the inverse filters applied. All the speakers were matched for level using a microphone positioned upright at the level of the ferret head in the centre of the semi-circle; correcting attenuations were applied to the stimuli before presentation.

In testing, stimuli were two 150 ms broadband noise bursts filtered according to the testing block being performed, including a 5 ms cosine envelope at onset and offset, presented sequentially from two speakers separated by 30° with a 20 ms silent gap between them. Locations tested were −75° to 75° at 30° intervals, in some sessions (BBN and BPN) speakers at −30°, 0° and 30° were added to the −75° to 75° speaker set. In a small number of early recordings (~3% of sessions from F1301 and F1302, for BBN 19/544 and LPN 11/339 recording sessions) speakers spanning −90° to 90° at 30° intervals were tested, in these recordings stimuli were also 200 ms duration. For these recordings spatial tuning properties and MI decoding was performed on the first 150 ms after stimulus onset. In population decoding only data from −75° to 75° at 30° intervals were evaluated.

### Frequency tuning protocol

To determine the Frequency Response Areas (FRAs) of any units being recorded from, ferrets were placed in an alternative testing arena with two speakers located on the left and right (24 cm from the ferret head) at head height of a central spout. Ferrets were provided with a constant stream of water from the central spout. The ferrets did not have to perform a task during an FRA recording session. Speakers were matched for level against each other and for presentation of all frequencies in an anechoic environment using a Brüel and Kjær 4191 condenser microphone attached to a Brüel and Kjær 3110-003 measuring amplifier. Sounds of varying frequency (150 Hz to 20 kHz at 1/3 octave intervals) at varying level (0 dB to 70 dB SPL) were presented to the ferrets while their head was at the central spout and recordings from auditory cortex were made.

The electrode arrays were constructed using Warp16 electrode arrays (Neuralynx Inc., Bozeman, MT), and comprised 16 (in a 4 x 4 configuration) individually moveable, high impedance (~2 MΩ), tungsten electrodes. Guide tubes into which electrodes were placed were approximately 800 μm apart from the centres.

Insulation was removed from the section of electrode that made contact with guide tube (~12 mm from the tip) using a hot soldering iron to melt the insulation and forceps to scrape the electrode clean. Electrodes were then inserted tail-end into the guide tube of the Warp16 under a microscope until the tip just disappeared from the end of the guide tube. The electrode was then moved a further 3 mm exposing the de-insulated part, and a 30° bend in the electrode was made before the electrode was trimmed approximately 3 mm from the bend and pushed back into the guide tube. All the guide tubes were filled with electrodes and the bottom of the Warp16 drive was then covered in Silastic (QWIK-SIL, WPI, Sarasota, FL) to prevent fluid wicking up the guide tubes after implantation, thus protecting the array from shorting. To prevent the Silastic from filling the guide tubes, a small drop of triple antibiotic ointment was applied to the end of each guide tube. Wires were soldered to the two ground contact points on the array. The base of the connector was strengthened with epoxy-resin (adapted from 56).

### Electrode implantation

Anaesthesia was induced by a single dose of a mixture of medetomidine (Domitor; 0.022 mg/kg/h; Pfizer) and ketamine (Ketaset; 5 mg/kg/h; Fort Dodge Animal Health). The ferret was intubated, placed on a ventilator (683 small animal ventilator; Harvard Apparatus) and ventilated with oxygen and isoflurane (1-3.5%) to maintain anaesthesia throughout the surgery. A local anaesthetic/analgesic (Marcaine, 0.5%) was injected under the skin where incisions were to be made. An incision was made along the midline of the ferret’s head and the connecting tissue cut to free the skin from the underlying muscle. The posterior 2/3 of the left temporal muscle was removed exposing the dorsal and lateral parts of the skull. Two anchor/ground screw holes were drilled into the posterior medial part of the skull and self-tapping bone screws inserted. A craniotomy was made over auditory cortex. The pre-assembled electrode array was put in place covering A1 using a micromanipulator so that the bottom of the array was in contact with the dura. The array was then retracted, the craniotomy filled with Silastic and the array replaced before the Silastic set. The ground wires of each implant were wound around each other and wound around the ground screws. The protective cap screw was secured in place around the array with dental cement. A metal bar with two nuts was placed in the centre of the head to provide a head-fixing device for later electrode movement. Further analgesia (Marcaine, 0.5%) was injected around the wound margin before the ferret was allowed to recover from the surgery. Post-operatively ferrets were given pain relief (buprenorphine, 0.01-0.03 mg/kg) for 3-5 days post-surgery and prophylactic antibiotics (Amoxycare LA, 15 mg/kg) and anti-inflammatories (Loxicam, 0.05 mg/kg) for 5 days post-surgery.

### Electrode moving

Electrodes were initially inserted into the brain approximately 1 week after surgery. The ferret was sedated with medetomodine (Domitor 0.022 mg/kg) and placed inside a plastic tube containing a heat pad. The temperature probe was placed between the ferret and the heat pad and the temperature was maintained at 38°C. The ferret’s head was held fixed using the nuts implanted during surgery. During the initial electrode insertion, the impedance of the electrodes was monitored (using an Omega-Z-Tip Impedance Meter, World Precision Instruments, UK) as they were pushed down into the brain using a Manual Cyborg Electrode Pusher (Neuralynx Inc., Bozeman, MT). The pusher probe guide tube was lowered around the electrode guide tube on the implant, and a probe wire was pushed down into the guide tube using a micro-manipulator (Harvard Instruments, USA). The electrode was pushed down until a drop in the impedance of the electrode was observed indicating the electrode had left the Silastic into which the array was implanted; the electrode was then pushed down a further 100-150 μm. All further movements were measured from this point which was defined as the surface of the brain. The electrodes were moved by 50-150 μm whenever the ferret had completed all required behavioural testing. In this manner over the course of 1-2 years recordings were made from each cortical layer in each ferret. The ferret completed the testing when the electrodes had moved a depth that exceeded the estimate for the maximal depth of auditory cortex (2 mm). The location and final depths of the electrodes were later confirmed by histology. These data, combined with estimates of frequency tuning made at each site, enabled an estimate of the location of each electrode in auditory cortex (Fig. 1).

### Spike sorting

The raw broadband voltage trace was filtered using an elliptical filter with bandwidth 300-5000 Hz (Matlab). The resulting filtered trace was processed to remove noise correlated across channels using methods described in Musial et al. (57). Spikes were detected using an amplitude threshold set automatically using methods described in Quiroga et al. (58). Spikes were clustered into “units” using algorithms adopted from Wave_Clus (58). Clusters were manually checked post-sorting and assigned as multi-unit or single-unit. Recording sessions performed at the same depth and within 3 days were combined and spike-sorted as if they were a single recording session. The spike shapes, rasters and PSTHs of these sessions were then checked manually for how well the recordings combined and were rejected if there was any inconsistency (e.g. if the spike shapes were different, if the PSTHs were different (T-test, p<0.05)), and were then spike sorted separately. Since multiple recordings were made at the same site in the brain, if the recordings could not be combined then the ‘best’ recording session for the particular site was taken (ensuring the same unit was not included in the analyses multiple times). The ‘best’ recording here refers either to the recording with the highest number of trials, or the recording with the best MI.

## Supplementary Information

**Table S1.** Behavioural performance statistics

| Stimulus | Ferret |  |  |  |
| --- | --- | --- | --- | --- |
|  | F1301 | F1302 | F1310 | F1313 |
| BBN | 865 / 1196<br>72.3%<br>$p < 0.001$ | 2726 / 3845<br>70.9%<br>$p < 0.001$ | 1557 / 2359<br>66.0%<br>$p < 0.001$ | 1563 / 2495<br>62.6%<br>$p < 0.001$ |
| LPN | 513 / 717<br>71.5%<br>$p < 0.001$ | 1566 / 2326<br>67.3%<br>$p < 0.001$ | 628 / 1005<br>62.5%<br>$p < 0.001$ | 468 / 800<br>58.5%<br>$p < 0.001$ |
| BPN | *Not tested* | 656 / 1106<br>59.3%<br>$p < 0.001$ | 580 / 991<br>58.5%<br>$p < 0.001$ | 459 / 810<br>56.7%<br>$p < 0.001$ |
| HPN | *Not tested* | 414 / 646<br>64.1%<br>$p < 0.001$ | 438 / 691<br>63.4%<br>$p < 0.001$ | 365 / 593<br>61.6%<br>$p < 0.001$ |
| CSS | *Not tested* | 795 / 1159<br>68.6%<br>$p < 0.001$ | 335 / 556<br>60.3%<br>$p < 0.001$ | 333 / 553<br>60.2%<br>$p < 0.001$ |

**Table S2.** Comparison of behaviour with limited cues or in the presence of a competing sound source with performance in broadband conditions

**F1301**
|  | Estimate | SE | t-statistic | p |
| --- | --- | --- | --- | --- |
| **intercept** | 0.961 | 0.065 | 14.863 | 0.000 |
| BBN:LPN | -0.038 | 0.105 | -0.366 | 0.714 |

Observations: 1913
Degrees of freedom: 1911

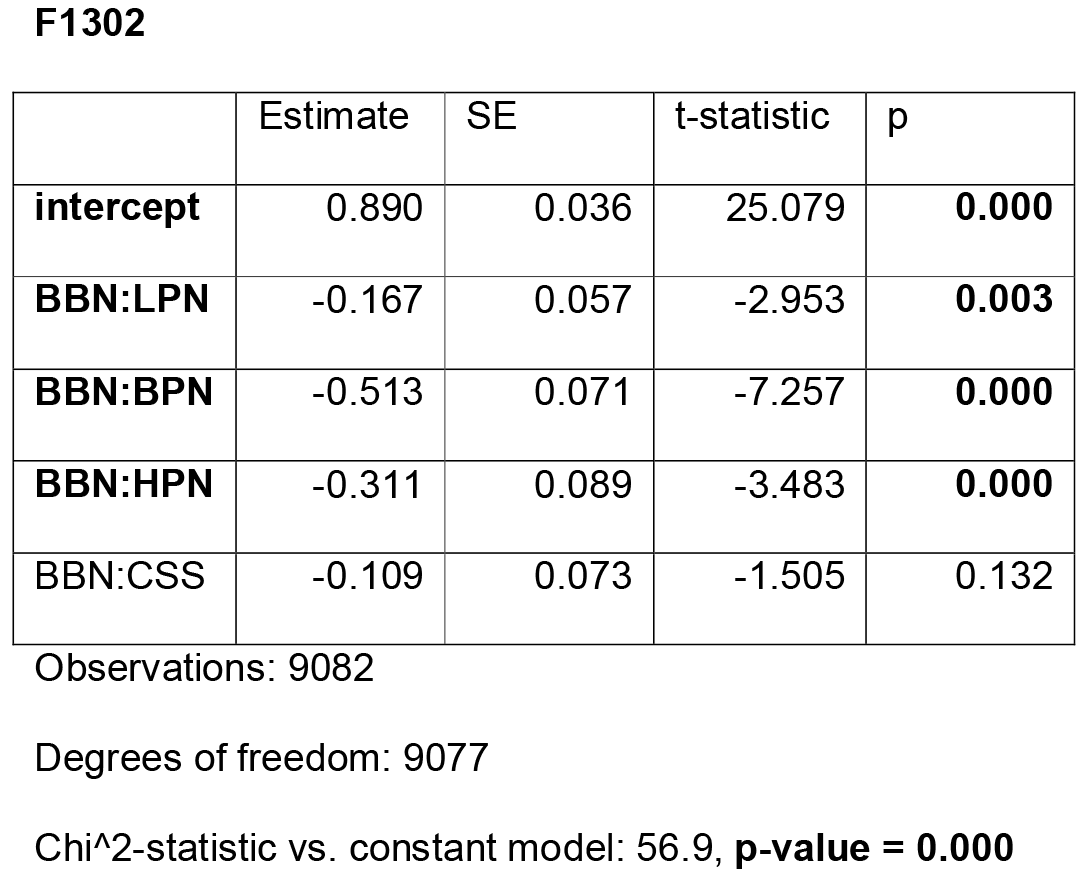

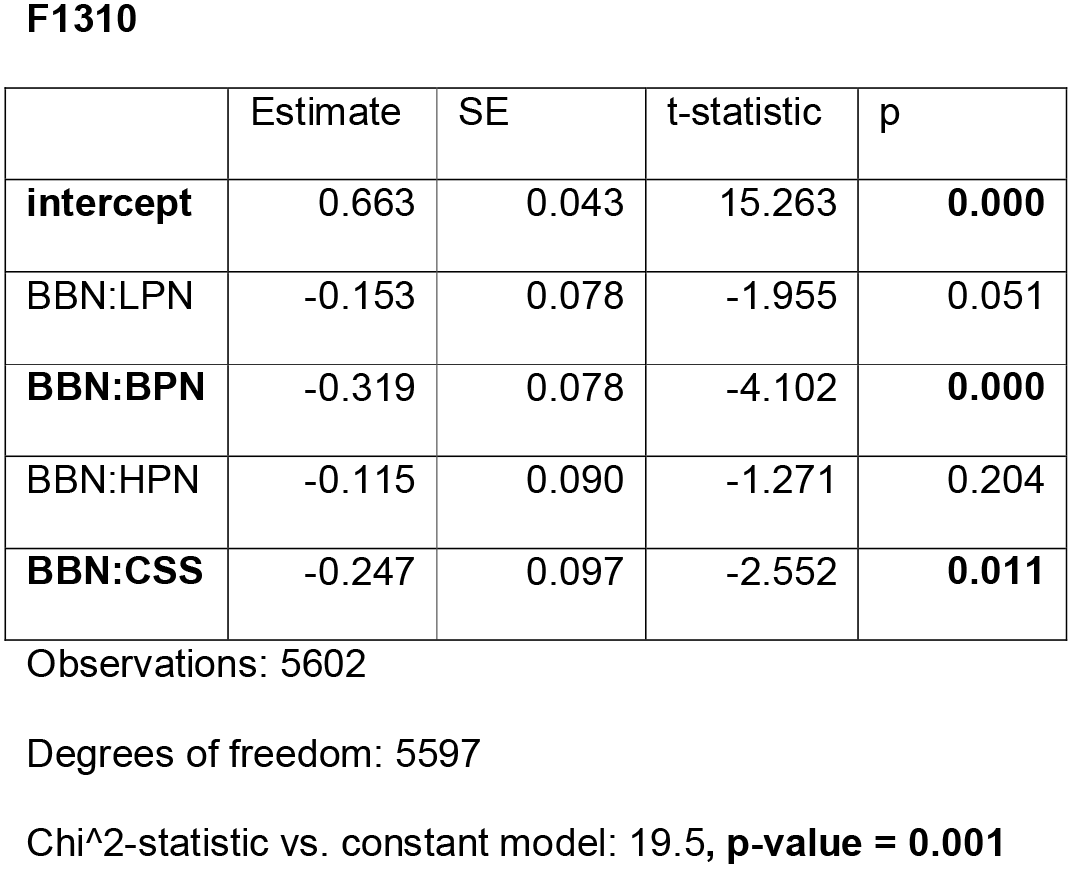

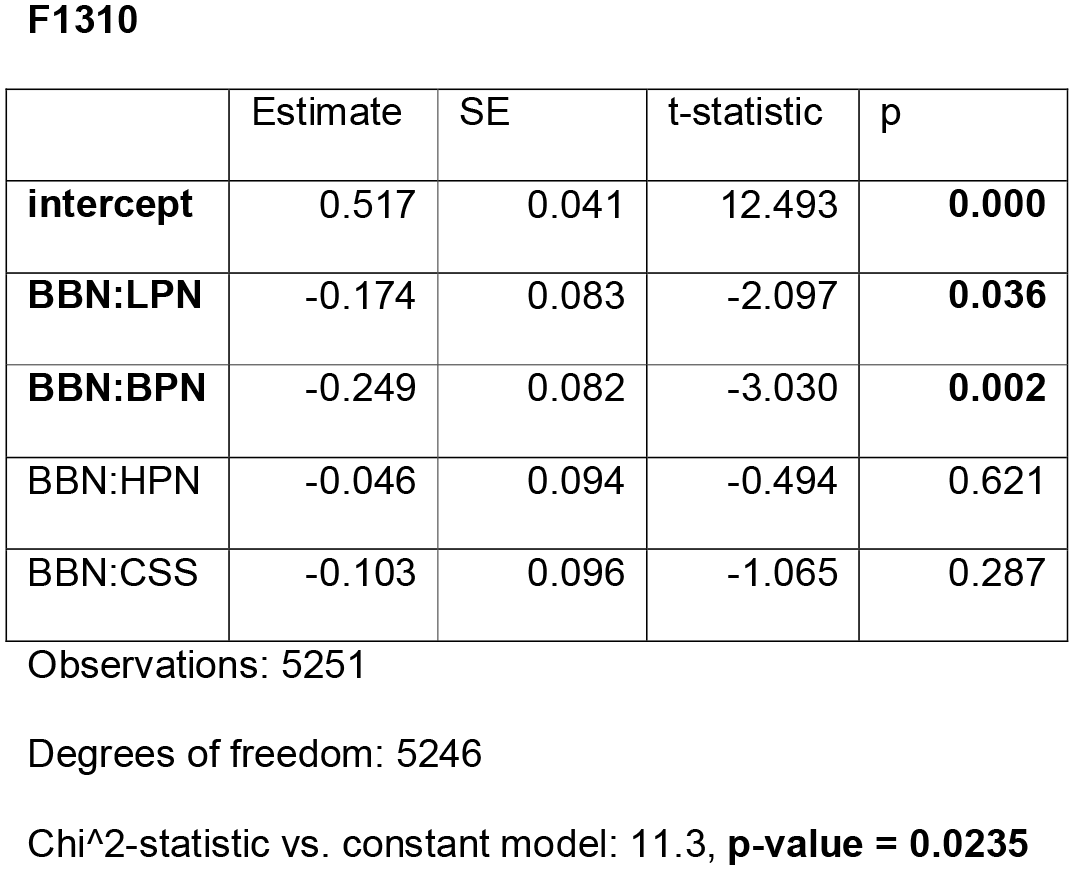

|  | Estimate | SE | t-statistic | p |
| --- | --- | --- | --- | --- |
| **intercept** | 0.890 | 0.036 | 25.079 | **0.000** |
| **BBN:LPN** | -0.167 | 0.057 | -2.953 | **0.003** |
| **BBN:BPN** | -0.513 | 0.071 | -7.257 | **0.000** |
| **BBN:HPN** | -0.311 | 0.089 | -3.483 | **0.000** |
| BBN:CSS | -0.109 | 0.073 | -1.505 | 0.132 |
| **intercept** | 0.663 | 0.043 | 15.263 | **0.000** |
| BBN:LPN | -0.153 | 0.078 | -1.955 | 0.051 |
| **BBN:BPN** | -0.319 | 0.078 | -4.102 | **0.000** |
| BBN:HPN | -0.115 | 0.090 | -1.271 | 0.204 |
| **BBN:CSS** | -0.247 | 0.097 | -2.552 | **0.011** |

**F1310**
|  | Estimate | SE | t-statistic | p |
| --- | --- | --- | --- | --- |
| **intercept** | 0.517 | 0.041 | 12.493 | **0.000** |
| **BBN:LPN** | -0.174 | 0.083 | -2.097 | **0.036** |
| **BBN:BPN** | -0.249 | 0.082 | -3.030 | **0.002** |
| BBN:HPN | -0.046 | 0.094 | -0.494 | 0.621 |
| BBN:CSS | -0.103 | 0.096 | -1.065 | 0.287 |

Observations: 5251
Degrees of freedom: 5246
Chi<sup>2</sup>-statistic vs. constant model: 11.3, **p-value = 0.0235**

**Fig. S1.**
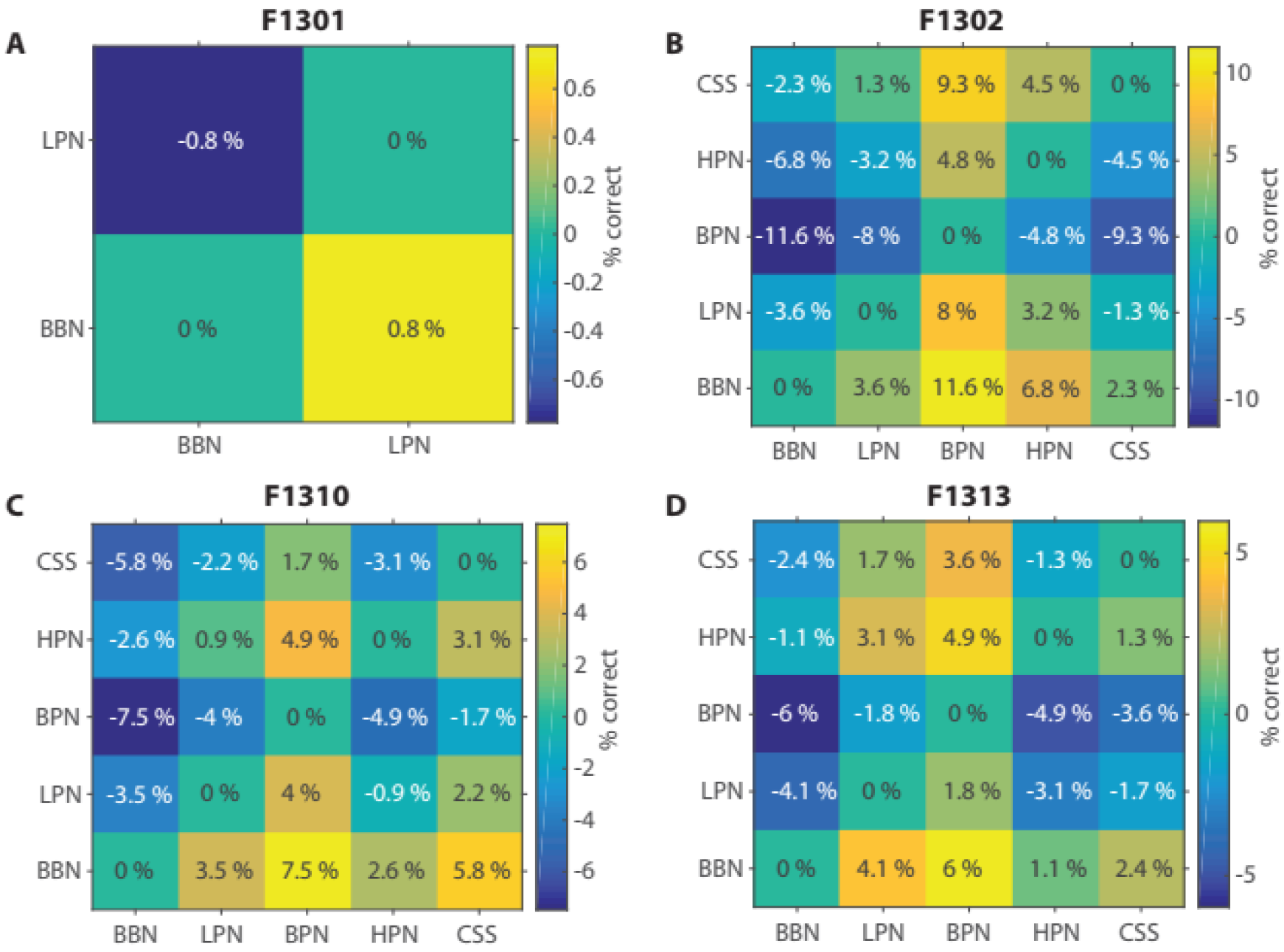
Change in performance between stimulus conditionsFig. S2 Comparison of bin width on decoder performance for broadband stimuli.

**Fig. S2.**
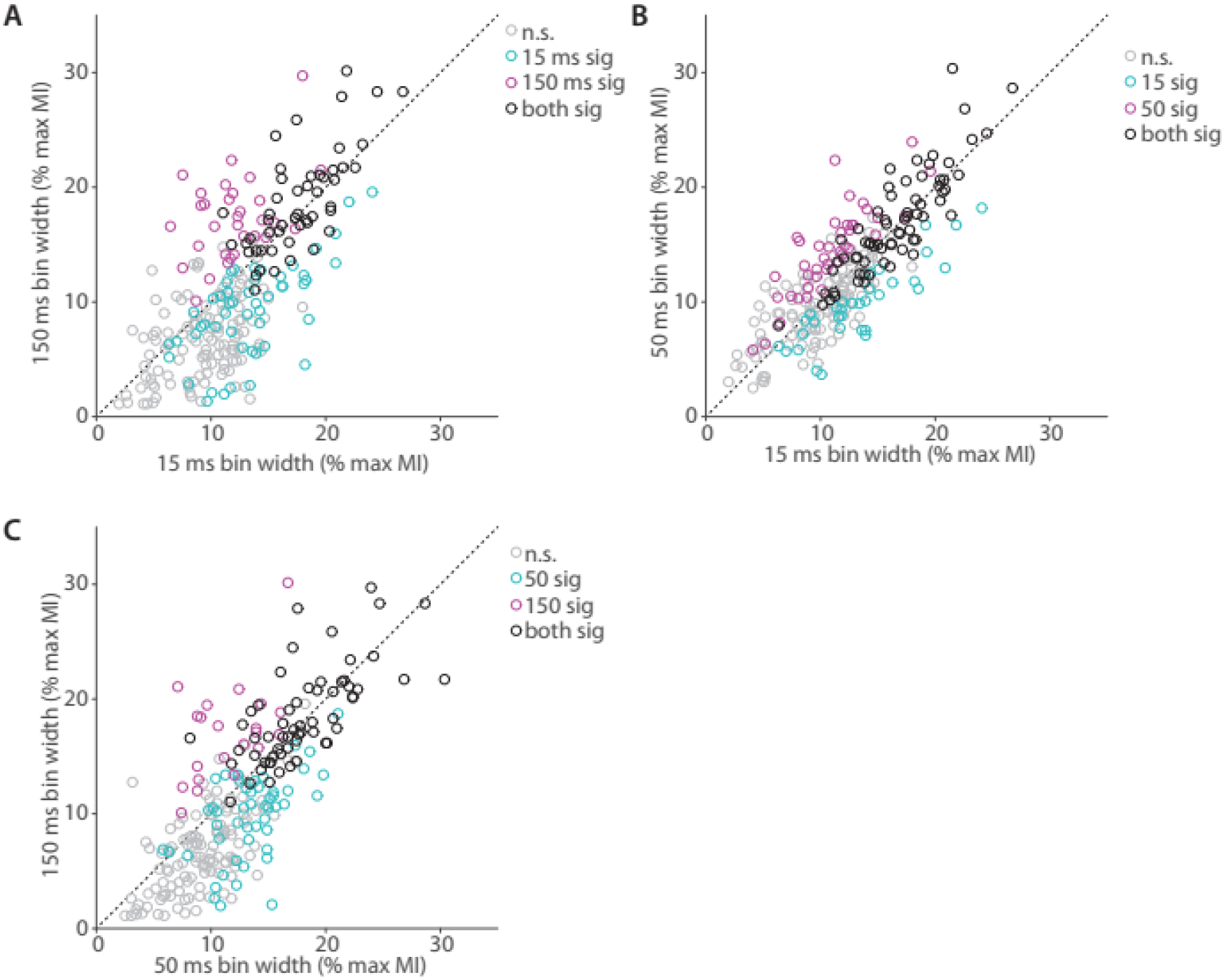
Comparison of bin width on decoder performance for broadband stimuli.

**Fig. S3.**
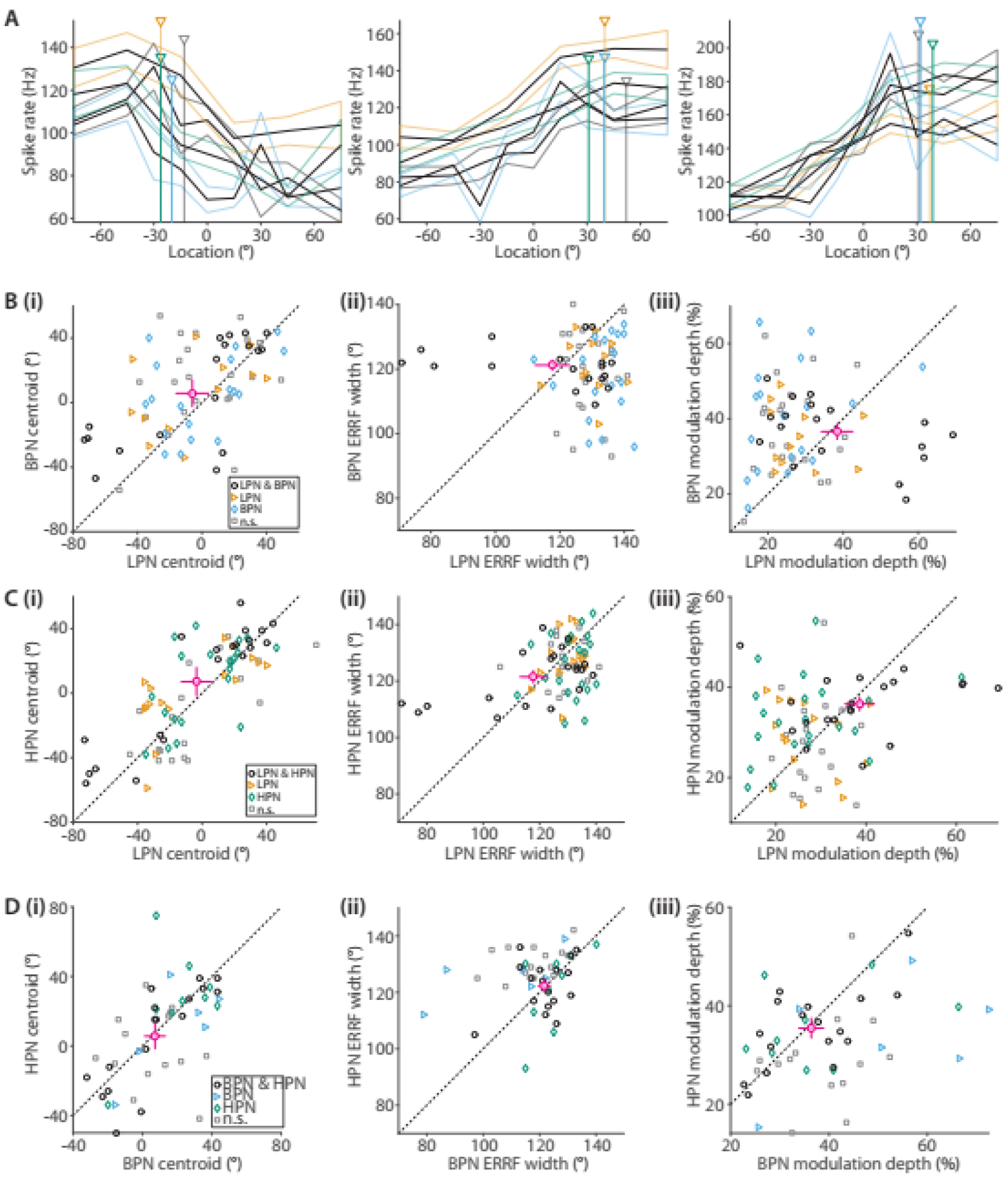
Comparison of spatial tuning properties of limited cue stimuli. (A) Shows three example units recorded in all 4 localisation cue conditions, broadband (BBN, grey), low-pass (LPN< orange), band-pass (BPN, blue) and high-pass (HPN, green). (B) Centroids (i), ERRF width (ii) and modulation depth of units recorded in both LPN and BPN conditions. Units that were spatially tuned in both conditions (circles), either condition alone (diamonds - BPN, triangles - LPN) or tuned in neither (squares). Mean ± standard error of the *paired units* mean of each is shown by crosshairs (circle, magenta). (C) Same as B for LPN and HPN stimuli. (D) Same as B for BPN and HPN stimuli

**Fig. S4.**
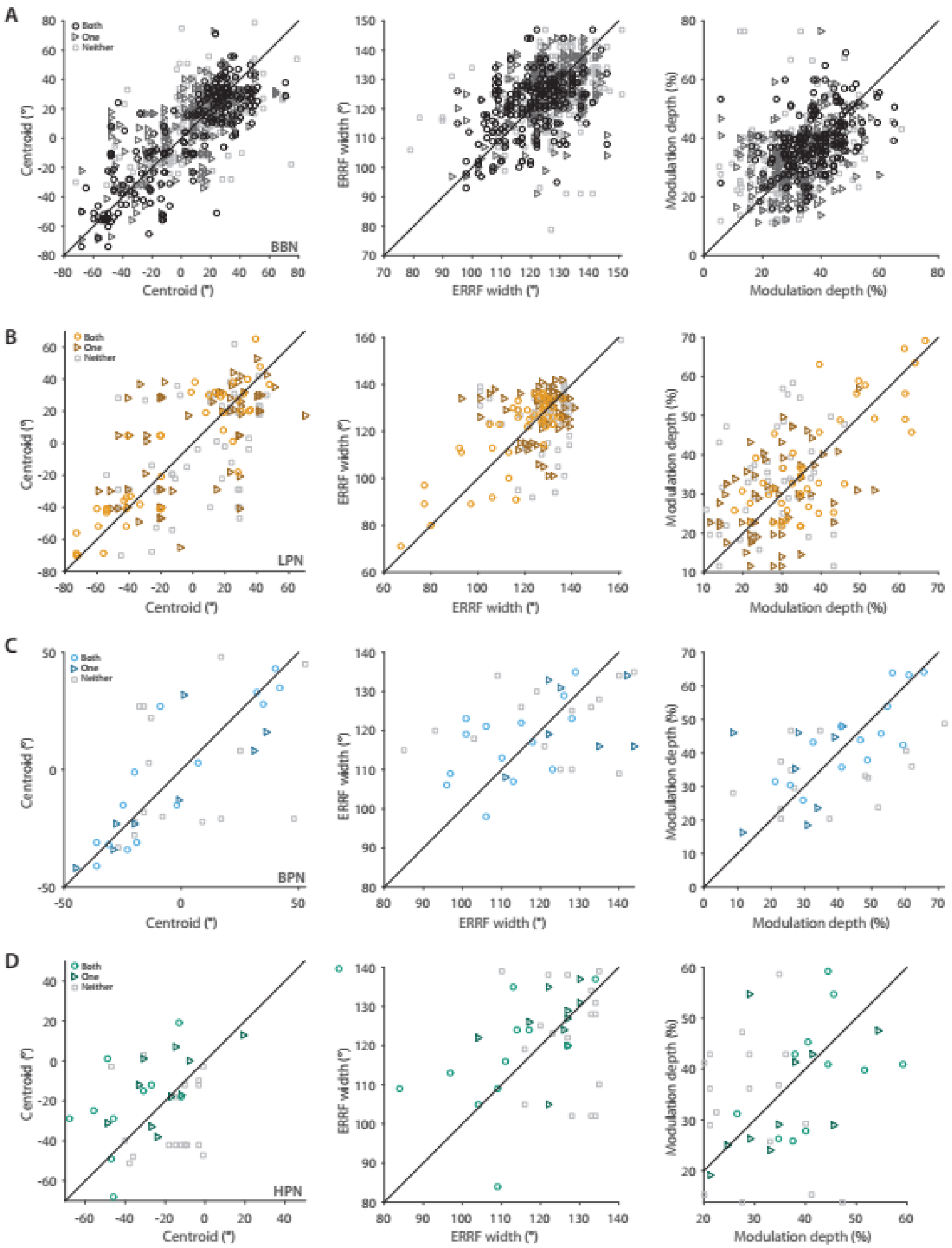
Comparison of centroid, ERRF width and modulation depth of units recorded at the same site. Comparison of centroids (1^st^ column), ERRF widths (2^nd^ column) and modulation depths (3^rd^ column) from units recorded at the same site in response to broadband (BBN, A), low-pass (LPN, B), band-pass (BPN, C) and high-pass (HPN, D) stimuli. Units were either both spatially modulated (circles), one of the pair spatially modulated (triangles) or neither spatially modulated (squares).

**Fig. S5.**
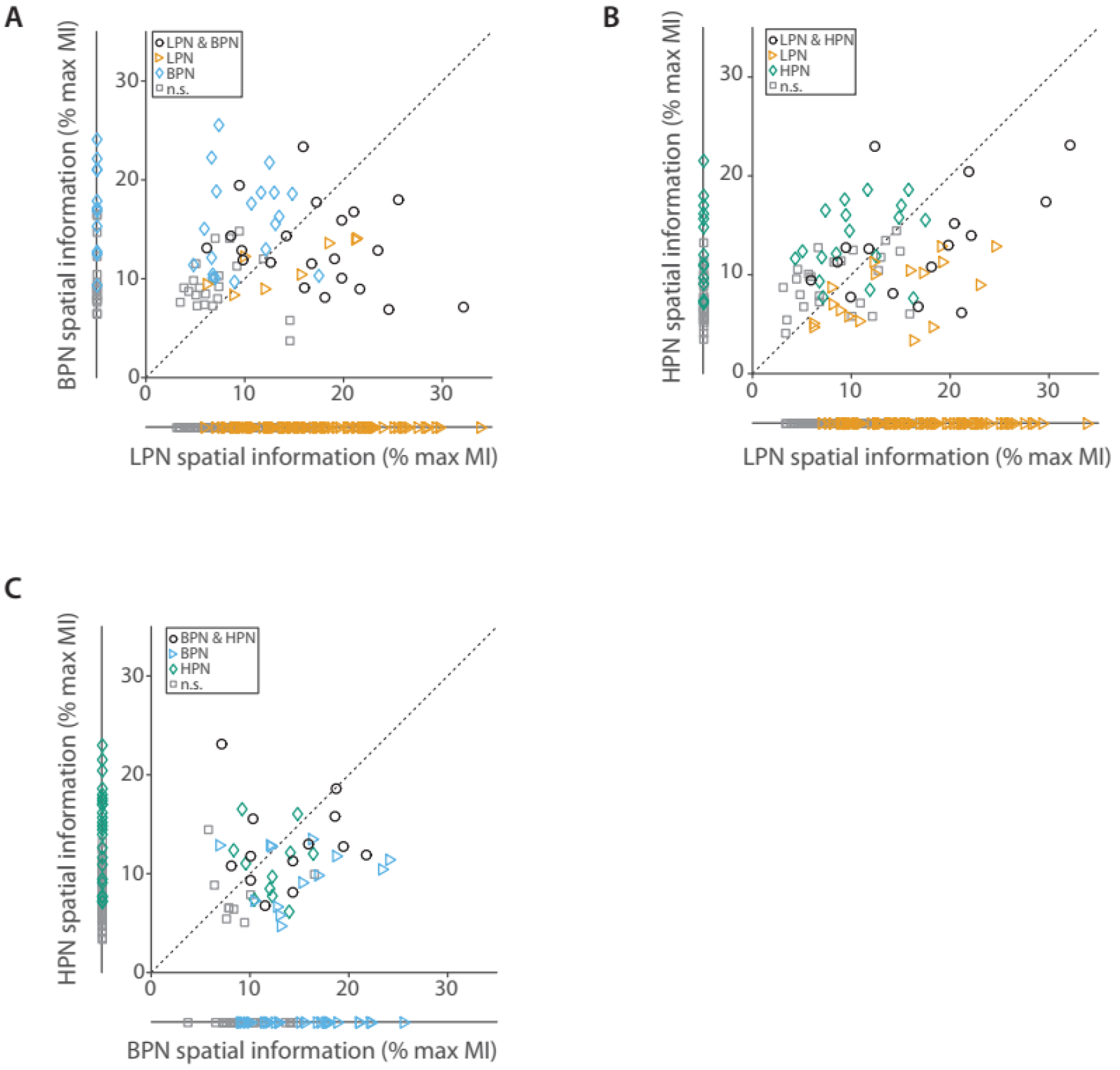
Comparison of spatial information between limited cue stimulus conditions.

**Fig. S6:**
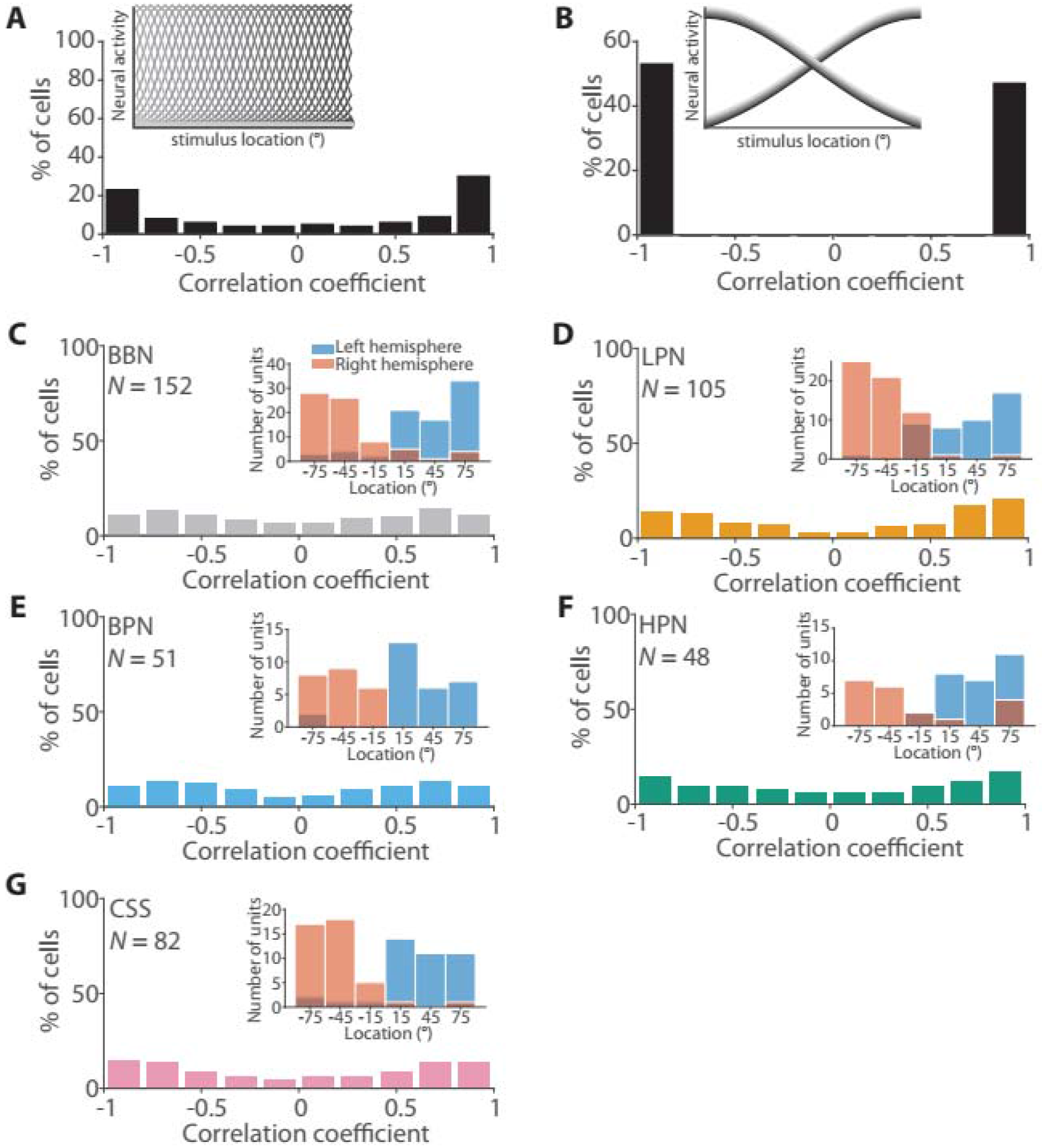
Tuning curve correlations resemble those of a labelled-line model. Distribution of correlation coefficients for modelled (inset) labelled line (A) and two-channel (B) tuning curves. (C-G) Distribution of correlation coefficients and best azimuths (inset) of tuning curves in each stimulus paradigm. Units had significant MI in at any bin width and were limited to those tested in the −75 to 75 testing locations so as to be comparable with performance in the population decoders (Fig. 7).

